# A polysynaptic hypothalamus–pelvic ganglion circuit regulates prostate tumor growth via WNT3A

**DOI:** 10.64898/2026.10.22.760909

**Authors:** Wang Ran, Peixuan Wu, Ziyun Qiao, Shuaidong Wang, Qiwei Yang, Rucheng Yan, Zhichao Yan, Jiajun Zhang, Biming He, Yongjia Shao, Yibin Wang, Shuguang Chu, Yongjie Wang, Haifeng Wang

**Affiliations:** Department of Urology, Shanghai East Hospital, School of Medicine, Tongji University, Shanghai, China; Department of Radiology, Shanghai East Hospital, Tongji University School of Medicine, Shanghai, China; School of Pharmacy, Hangzhou Normal University, Hangzhou, China

**Keywords:** Prostate cancer, Brain – tumor communication, Polysynaptic neural circuits, Autonomic innervation, Major pelvic ganglion, WNT3A

## Abstract

Brain – tumor communication is increasingly recognized. Peripheral autonomic nerves regulate prostate cancer progression, yet how the brain influences local nerve–tumor interactions remains unclear. We identify a polysynaptic circuit linking the hypothalamic paraventricular nucleus (PVN), dorsal motor nucleus of the vagus (DMV), major pelvic ganglion (MPG) and prostate tumor in mice. Although both nuclei were activated during tumor growth, PVN activity restrained, whereas DMV activity promoted tumor growth. MPG ablation abolished DMV-induced tumor promotion. Combining LMD with Smart-seq2 revealed transcriptional remodeling and increased Wnt3a expression in tumor-innervating, activated MPG neurons under upstream neural control. MPG explants released WNT3A, whose neutralization suppressed explant-induced cancer-cell proliferation. Together, our findings identify a polysynaptic PVN–DMV–MPG circuit that regulates prostate tumor growth and implicate WNT3A as a mediator of the pro-tumor effects of the MPG, highlighting the therapeutic potential of targeting the neural circuit and downstream molecular pathway.

## Introduction

Prostate cancer is the second most commonly diagnosed malignancy and the fifth leading cause of cancer-related death among men worldwide^1–3^. During tumor progression, cancer cells can hijack neural regulatory mechanisms to acquire phenotypes that support growth and invasion^4^. The prostate is richly innervated, and previous studies have shown that both sympathetic and parasympathetic signaling can promote prostate cancer progression through mechanisms involving angiogenesis, metabolic regulation and epigenetic reprogramming^5–8^.

However, most studies of neural–tumor interactions have focused on nerve terminals within the tumor, whereas the neuronal cell bodies that contain the relevant transcriptional information reside in peripheral ganglia outside the tumor^9,10^. Whether neurons that directly innervate prostate tumor undergo specific molecular remodeling, and whether such changes are regulated by upstream neural circuits, remain poorly understood. The major pelvic ganglion (MPG) is a key autonomic ganglion innervating pelvic organs and contributes to urinary, gastrointestinal and reproductive functions. It receives sympathetic preganglionic input through the hypogastric nerve and parasympathetic preganglionic input through the pelvic nerve, making it a mixed autonomic ganglion^11,12^. However, the role of the MPG in prostate cancer progression and the underlying molecular mechanisms remain unclear.

Peripheral nerves that innervate tumor are regulated by the central nervous system. The hypothalamic paraventricular nucleus (PVN) is a major integrative centre for sympathetic and parasympathetic activity^13^. Beyond its classical neuroendocrine functions, parvocellular preautonomic neurons in the PVN project broadly to autonomic nuclei in the brainstem and spinal cord, including the nucleus of the solitary tract, dorsal motor nucleus of the vagus (DMV) and rostral ventrolateral medulla^14^. Recent studies have further shown that peripheral tumor can increase neuronal activity in several brain regions, including the PVN, and that distinct PVN neuronal populations regulate lung, breast and colorectal cancer progression through neuroendocrine and autonomic pathways^15–18^. However, whether the PVN is connected to prostate tumor through a multisynaptic circuit, and how this circuit regulates tumor progression, remain largely unknown.

Here, we identified anatomical and functional evidence supporting a PVN–DMV–MPG-associated multisynaptic pathway to prostate tumor and examined how manipulation of these nodes affects tumor growth. By combining laser microdissection (LMD) with Smart-seq2^19,20^, we characterized the transcriptional profiles of activated MPG neurons that innervate prostate tumor and assessed how PVN activation reshaped their molecular state. We further found that WNT3A neutralization suppressed MPG-associated prostate tumor cell proliferation and invasion. Together, our findings identify the PVN and DMV as key central autonomic nodes in prostate cancer-associated neural circuitry and highlight neural communication between the brain and tumor as a potential therapeutic target.

## Results

### Tumor-innervating MPG neurons are activated in prostate tumor-bearing mice

In the orthotopic prostate tumor model, sympathetic and cholinergic nerve fibers were detected in both normal prostate and tumor tissues, with increased neural infiltration at the tumor margins. In Ai14/FOS-CreERT2 mice with prostate tumor, in which Fos-expressing neurons were permanently labeled with tdTomato, axon terminals of tdTomato⁺ FOS activated neurons were also detected within the tumors (Extended Data Fig. 1A–C).

Consistent with these findings, TCGA analysis identified marked upregulation of the neural-associated gene DLX1 in prostate cancer (Extended Data Fig. 1D). Single-cell RNA sequencing of prostate cancer biopsies from 6 patients showed higher PNI and neuronal marker scores in malignant epithelial cells than in normal epithelial cells, with C3, C6 and C7 displaying prominent neural-related transcriptional features (Extended Data Fig. 1E–I). Among the significantly upregulated genes in the PNI-marker, neural-marker and composite-neural-score gene sets, high expression of CLDN3, MDK, ARHGDIB and PLK2 was associated with an increased risk of biochemical recurrence in the TCGA cohort (Extended Data Fig. 1J), further linking neural-related molecular features to prostate cancer progression.

To extend these clinical observations, we next examined neuronal activation and prostate tumor innervation across peripheral ganglia in mice. FOS-activated neurons were detected in the MPG and S1 DRG of Ai14/FOS-CreERT2 mice with prostate tumor (Fig. 1A–B). The proportion of activated neurons in the MPG was significantly higher in tumor mice than in control or benign prostatic hyperplasia mice (Fig. 1B, all P < 0.05). DiI retrograde tracing identified tumor-innervating neurons in the DRG, MPG and IMG, with preferential enrichment in the S1 DRG and MPG (Fig. 1C–E). TUBB3 staining confirmed the neuronal characteristics of these ganglia, and CTB tracing independently supported MPG innervation of prostate tumor (Extended Data Fig. 2A–C). Furthermore, PRV-EGFP tracing combined with FOS-dependent labelling identified PRV⁺/FOS⁺ neurons in the MPG (Fig. 1F–H), indicating the presence of MPG neurons that both innervate the prostate tumor and are simultaneously activated. Immunofluorescence staining showed that these activated MPG neurons were ChAT⁺ but TH⁻ (Extended Data Fig. 2D), supporting preferential activation of the cholinergic parasympathetic component during prostate tumor progression.

**Fig. 1.**
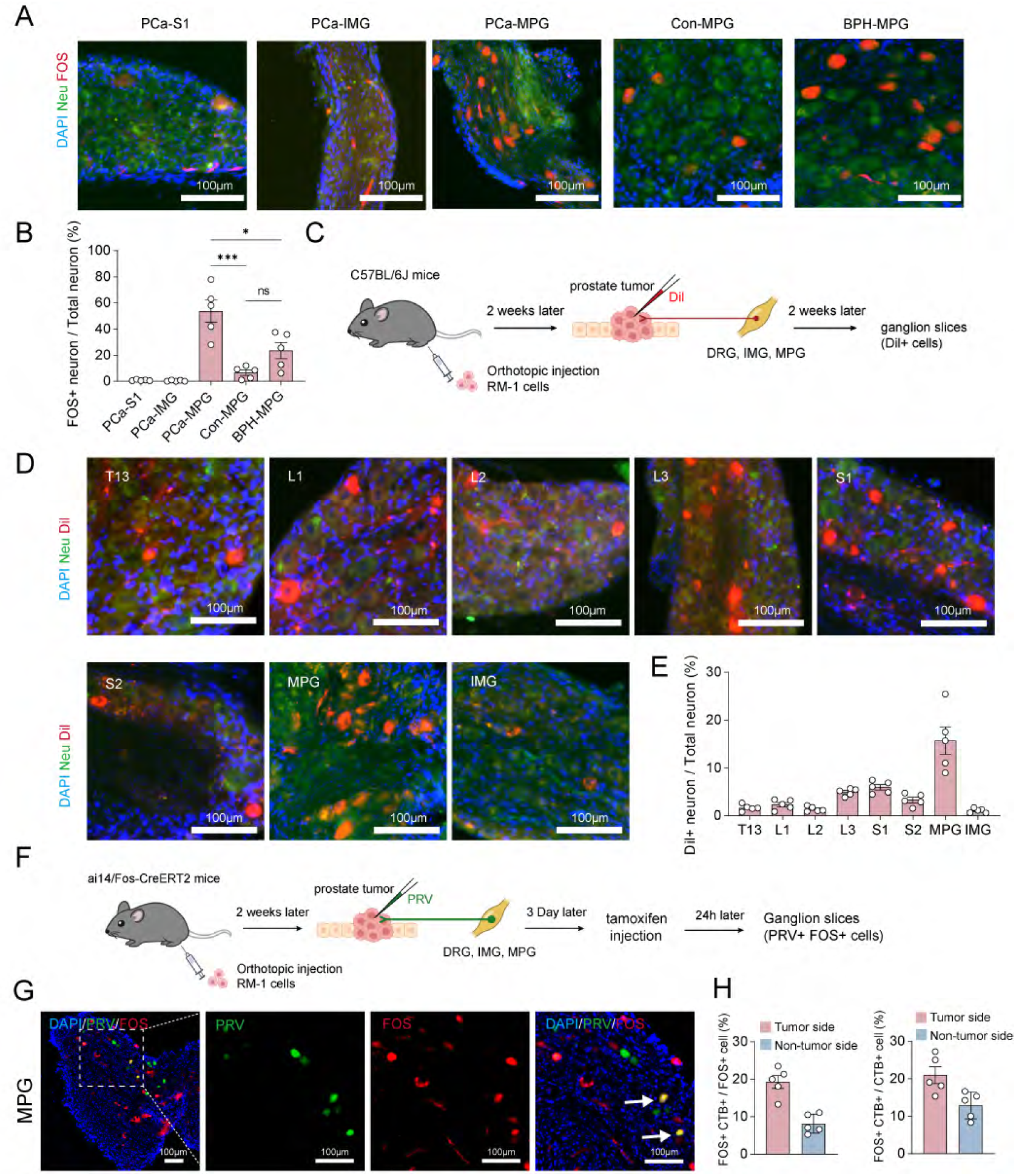
MPG innervation and activation in prostate tumor. **A**, Representative immunofluorescence images showing neuronal FOS expression in the S1 DRG, IMG and MPG of PCa mice and in the MPG of Con and BPH mice. Neuronal somata, shown in the green channel, were identified based on typical ganglionic morphology and intrinsic autofluorescence in the 488-nm channel. DRG, dorsal root ganglion; IMG, inferior mesenteric ganglion; MPG, major pelvic ganglion; PCa prostate cancer; Con, control; BPH, benign prostatic hyperplasia. **B,** The proportion of FOS⁺ neurons in the MPG was significantly higher in the PCa group than in the control and BPH groups. n = 5 mice. One-way ANOVA followed by Tukey’s multiple-comparisons test. *P < 0.05, ***P < 0.001; ns, not significant. **C,** Schematic of DiI retrograde tracing from orthotopic RM-1 prostate tumors, followed by analysis of DiI⁺ neurons in the DRGs, IMG and MPG. **D**, Representative fluorescence images showing Dil-labelled prostate tumor-innervating neurons in the T13, L1, L2, L3, S1 and S2 DRGs, MPG and IMG. **E,** Quantification of the proportion of Dil⁺ neurons among total neurons in the indicated ganglia. n = 5 mice. **F,** Schematic of PRV tracing and activity-dependent neuronal labeling in orthotopic RM-1 tumors in Ai14/FOS-CreERT2 mice, followed by analysis of PRV⁺/FOS⁺ neurons in peripheral ganglia. **G**, Representative fluorescence images showing PRV-labelled and FOS-activated neurons in the MPG of tumor mice. **H,** Quantification of PRV⁺/FOS⁺ double-positive neurons as a percentage of total FOS⁺ neurons (left) or total PRV⁺ neurons (right) in the tumor-side and non-tumor-side MPG. n = 5 mice.

### Tumor-connected PVN and DMV neurons are activated in prostate tumor-bearing mice

Previous studies have shown that the central nervous system responds to peripheral tumor growth, yet the brain circuits associated with prostate cancer remain poorly defined. We therefore sought to identify central neural nodes upstream of the MPG.

In Ai14/FOS-CreERT2 mice with prostate tumor, FOS⁺ neurons were significantly increased in several brain regions, including PVN, DMV, paraventricular nucleus of the thalamus (PV), subfornical organ (SFO), parataenial nucleus (PT), suprachiasmatic nucleus (SCh), arcuate nucleus (ARC) and median eminence (ME) (all P < 0.05; Extended Data Fig. 3A–B and Fig. 2A). Trans-synaptic retrograde tracing from the prostate with PRV-EGFP labelled neurons in the PVN, DMV, medial preoptic area (MPO), red nucleus (RN), periaqueductal grey (PAG), pontine micturition centre (PMC), locus coeruleus (LC), gigantocellular reticular nucleus (Gi) and raphe obscurus nucleus (ROb), indicating multisynaptic connectivity between the tumor and multiple central nuclei (Extended Data Fig. 3C and Fig. 2A). Combining PRV-EGFP tracing with FOS-dependent labelling further identified PRV⁺/FOS⁺ neurons in the PVN and DMV (Fig. 2B), demonstrating that these neurons were both connected to the prostate tumor-associated circuit and activated in tumor mice. Most activated PVN neurons were OXT⁺, whereas activated DMV neurons expressed OXTR and were predominantly cholinergic (Fig. 2C–D).

**Fig. 2.**
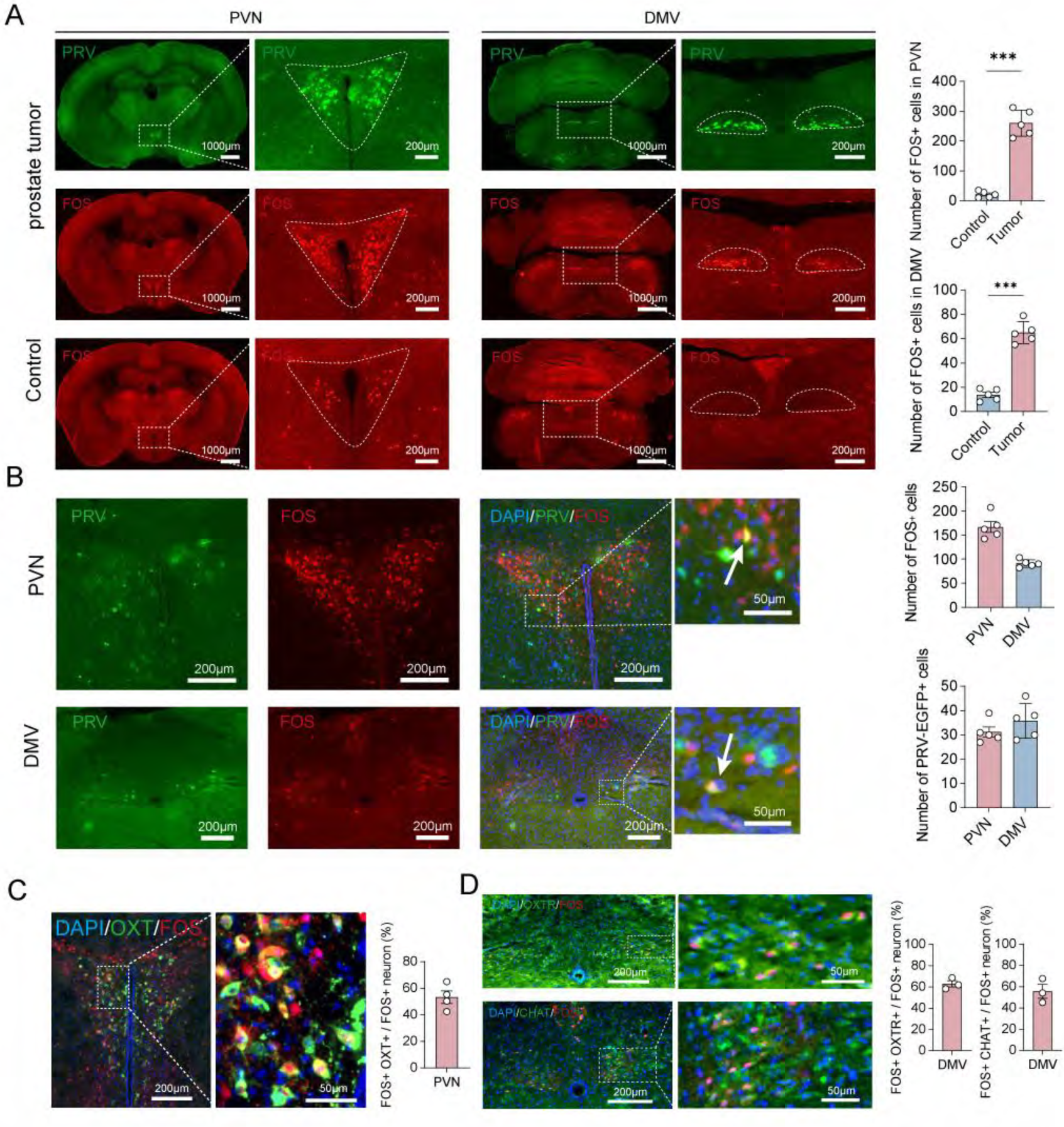
PVN and DMV innervation and activation in prostate tumor. **A**, Representative fluorescence images showing PRV-labelled neurons and FOS activation in the PVN and DMV in prostate tumor-bearing mice and healthy control mice. Enlarged images of the indicated regions are shown on the right. Quantification showed significantly increased numbers of FOS⁺ cells in the PVN and DMV of tumor-bearing mice compared with control mice. n = 5 mice per group. Two-tailed unpaired Student’s t-test. ***P < 0.001; ****P < 0.0001. **B,** Representative fluorescence images showing PRV-labelled and FOS-activated neurons in the PVN and DMV of prostate tumor-bearing mice. Enlarged images show PRV⁺/FOS⁺ double-positive neurons (white arrows). Quantification of the numbers of FOS⁺ cells and PRV/EGFP⁺ cells in the PVN and DMV is shown on the right. n = 5 mice per group. **C,** Representative immunofluorescence images showing colocalization of OXT with FOS-activated neurons in the PVN. The graph shows the proportion of FOS⁺/OXT⁺ double-positive neurons among FOS⁺ neurons in the PVN. n = 3 mice. OXT, oxytocin. **D,** Representative immunofluorescence images showing colocalization of OXTR or ChAT with FOS-activated neurons in the DMV. Quantification of FOS⁺/OXTR⁺ and FOS⁺/ChAT⁺ double-positive neurons among FOS⁺ neurons are shown on the right. n = 3 mice. OXTR, oxytocin receptor.

### Anatomical tracing supports a PVN–DMV–MPG-associated multisynaptic pathway to prostate tumor

To define the temporal sequence of trans-synaptic labelling, PRV-EGFP was injected into prostate tumor of C57BL/6J mice, and neural tissues were collected at 72, 96 and 120 h after injection (Fig. 3A). At 72 h, PRV⁺ neurons were detected in MPG, whereas no labelling was observed in the brain. PRV⁺ neurons subsequently appeared in DMV at 96 h and in PVN at 120 h (Fig. 3B). This sequential labelling pattern was consistent with a multisynaptic pathway linking the PVN, DMV, MPG and prostate tumor.

**Fig. 3.**
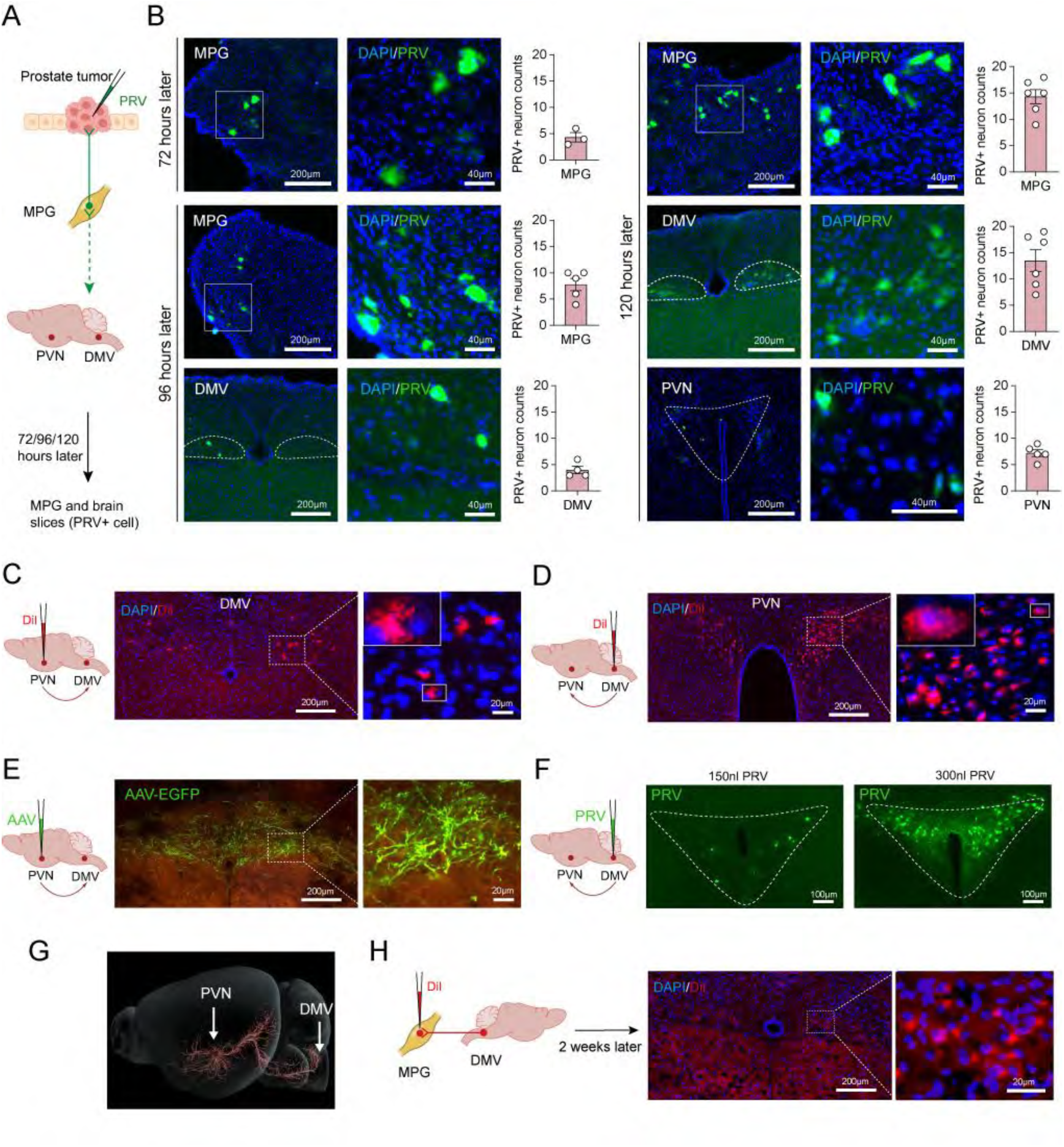
Neural circuit linking brain regions and the MPG to prostate tumor. **A**, Schematic of the experimental design for time-dependent trans-synaptic retrograde tracing from prostate tumors. **B,** Representative fluorescence images and quantification of PRV⁺ neurons in the MPG, DMV and PVN at 72, 96 and 120 h after intratumoral PRV-EGFP injection. PRV⁺ neurons were detected in the MPG at 72 h, subsequently in the DMV at 96 h and in the PVN at 120 h. n = 3, 4, 5 and 6 mice for the respective groups. **C,** Schematic and representative fluorescence images showing DiI-labelled neurons in the DMV following Dil injection into the PVN, consistent with anatomical connectivity between the PVN and DMV. **D,** Schematic and representative fluorescence images showing Dil-labelled neurons in the PVN following Dil injection into the DMV, further supporting anatomical connectivity between the PVN and DMV. **E**, Representative fluorescence images from an anterograde tracing dataset in the Allen Mouse Brain Connectivity Atlas, experiment 299759881. Cre-dependent AAV-EGFP was injected into the PVN of Efr3a-Cre mice to label neuronal projections, revealing EGFP-labelled axonal fibres in the DMV. **F,** Representative fluorescence images showing PRV-labelled neurons in the PVN following PRV injection into the DMV. Images show PVN labelling after injection of 150 nl or 300 nl PRV into the DMV. **G,** Three-dimensional reconstruction of the PVN-to-brainstem projection based on the Allen Mouse Brain Connectivity Atlas dataset, experiment 299759881, showing descending axonal fibers from the PVN traversing or terminating within the DMV region. **H**, Schematic and representative fluorescence images showing Dil-labelled neurons in the DMV 2 weeks after Dil injection into the MPG, supporting anatomical connectivity between the DMV and MPG.

To assess anatomical connectivity between the PVN and DMV, we performed non-trans-synaptic tracing with DiI. Injection of DiI into the PVN produced positive labelling in the DMV, whereas injection into the DMV labelled the PVN (Fig. 3C–D), supporting anatomical connectivity between these regions. Analysis of experiment 299759881 from the Allen Mouse Brain Connectivity Atlas further showed that injection of the Cre-dependent AAV-EGFP into the PVN of Efr3a-Cre mice labelled axonal fibres within the DMV, consistent with the direct projection from PVN to DMV (Fig. 3E). Moreover, PRV injection into the DMV labelled PVN neurons as early as 36 h, when extensive trans-synaptic spread may not yet have occurred (Fig. 3F). Three-dimensional reconstruction of the Allen dataset further showed that most descending fibers from the PVN to the brainstem traversed or terminated within the DMV region (Fig. 3G). Finally, DiI injection into the MPG resulted in positive labelling in the DMV after 2 weeks (Fig. 3H), supporting an anatomical connection between the DMV and MPG. Together, these findings identify the PVN–DMV–MPG axis as a potential anatomical substrate of the autonomic circuit associated with prostate tumor.

### Chemogenetic manipulation of tumor-activated PVN and DMV neurons regulates prostate tumor growth

To selectively inhibit tumor-activated neurons in PVN, AAV-DIO-hM4Di-EGFP or the control virus AAV-DIO-EGFP was injected into the PVN of FOS-CreERT2 mice with prostate tumor (Fig. 4A). Administration of clozapine-N-oxide (CNO) reduced FOS expression in hM4Di-expressing EGFP⁺ neurons, confirming effective neuronal inhibition (Fig. 4B). Inhibition of tumor-activated PVN neurons significantly increased prostate tumor growth (Fig. 4C–D). Consistently, selective ablation of these neurons using AAV-FLEX-taCasp3 also increased tumor growth (Extended Data Fig. 4A–D). Conversely, CNO increased FOS expression of hM3Dq-expressing EGFP⁺ neurons in the PVN (Fig. 4E), and chemogenetic activation of these neurons significantly suppressed prostate tumor growth (Fig. 4F–G).

**Fig. 4.**
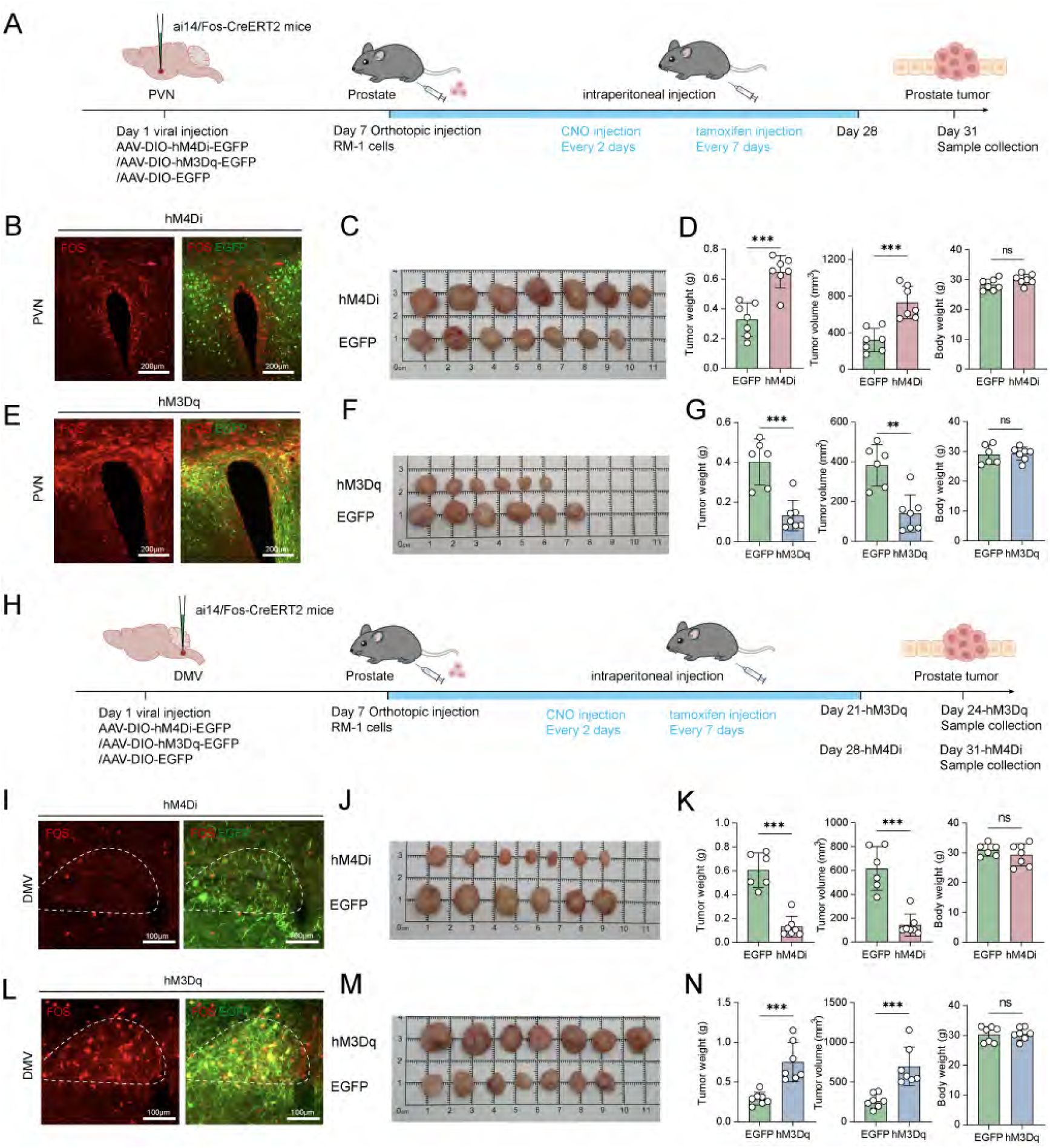
Interventions targeting activated PVN and DMV neurons affect prostate tumor growth. **A**, Experimental scheme for chemogenetic manipulation of tumor-activated PVN neurons. Ai14/FOS-CreERT2 mice received PVN injections of AAV-DIO-hM4Di-EGFP, AAV-DIO-hM3Dq-EGFP or control AAV-DIO-EGFP. Seven days later, RM-1 cells were orthotopically implanted into the prostate, followed by CNO and tamoxifen administration according to the indicated schedule. **B–D,** Chemogenetic inhibition of tumor-activated PVN neurons. Representative fluorescence images of FOS expression in hM4Di-EGFP⁺ PVN neurons (B), macroscopic images of prostate tumors (C) and quantification of tumor weight, tumor volume and body weight (D). PVN inhibition increased tumor weight and volume without affecting body weight. n = 7 mice per group. **E–G,** Chemogenetic activation of tumor-activated PVN neurons. Representative fluorescence images of FOS expression in hM3Dq-EGFP⁺ PVN neurons (E), macroscopic images of prostate tumors (F) and quantification of tumor weight, tumor volume and body weight (G). PVN activation reduced tumor weight and volume without affecting body weight. n = 6 mice per group. **H,** Experimental scheme for chemogenetic manipulation of tumor-activated DMV neurons using the same viral, tumor implantation and drug administration strategy. **I–K,** Chemogenetic inhibition of tumor-activated DMV neurons. Representative fluorescence images of FOS expression in hM4Di-EGFP⁺ DMV neurons (I), macroscopic images of prostate tumors (J) and quantification of tumor weight, tumor volume and body weight (K). DMV inhibition reduced tumor weight and volume without affecting body weight. Control, n = 6 mice; hM4Di, n = 7 mice. **L–N,** Chemogenetic activation of tumor-activated DMV neurons. Representative fluorescence images of FOS expression in hM3Dq-EGFP⁺ DMV neurons (L), macroscopic images of prostate tumors (M) and quantification of tumor weight, tumor volume and body weight (N). DMV activation increased tumor weight and volume without affecting body weight. n = 7 mice per group. Data are presented as mean ± SD. Two-tailed unpaired Student’s t-tests were used. **P < 0.01, ***P < 0.001; ns, not significant.

The same chemogenetic strategy was applied to tumor-activated neurons in the DMV (Fig. 4H). In contrast to the effects observed in the PVN, inhibition of DMV neurons significantly reduced prostate tumor growth (Fig. 4I–K), whereas their activation significantly promoted tumor growth (Fig. 4L–N). Consistently, selective ablation of these neurons using AAV-FLEX-taCasp3 also reduced tumor growth (Extended Data Fig. 4E–H). Together, these findings indicate that tumor-activated PVN and DMV neurons exert opposing effects on prostate tumor progression.

### PVN and DMV regulation of prostate tumor growth mediated by the MPG

Given the opposing effects of PVN and DMV manipulation on prostate tumor progression, we next assessed how these interventions altered neuronal activity at downstream nodes of the circuit. In tumor mice, hM3Dq-mediated activation of tumor-responsive PVN neurons significantly reduced FOS expression in the DMV, whereas hM4Di-mediated inhibition increased DMV FOS expression (Fig. 5A–B). Conversely, activation of tumor-responsive DMV neurons significantly increased FOS expression in MPG neurons, whereas DMV inhibition reduced it (Fig. 5C–D). PVN manipulation also affected MPG neuronal activity, with PVN activation reducing FOS expression in MPG neurons and PVN inhibition increasing it (Fig. 5E–F). These findings suggest that the PVN suppresses the prostate tumor-associated autonomic circuit by restraining DMV–MPG activity.

**Fig. 5.**
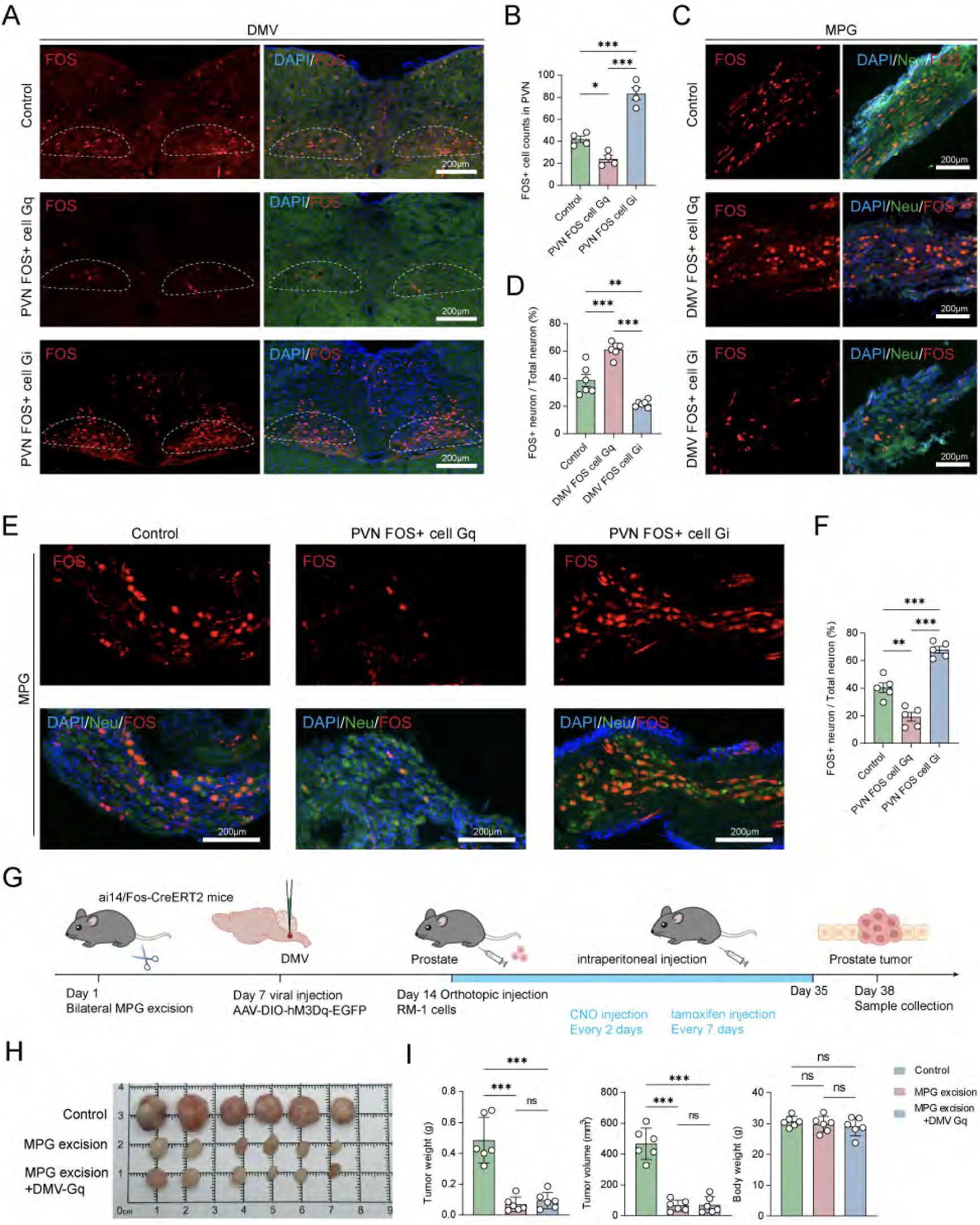
PVN and DMV regulation of MPG neuronal activity and attenuation of DMV-driven tumor growth by MPG excision. **A,B**, Representative fluorescence images (A) and quantification (B) of FOS activation in the DMV following chemogenetic manipulation of tumor-responsive neurons in the PVN. Activation of PVN neurons with hM3Dq reduced the number of FOS⁺ cells in the DMV, whereas inhibition with hM4Di increased DMV neuronal activation. Dashed lines delineate the DMV. n = 4 mice. **C,D,** Representative fluorescence images (C) and quantification (D) of FOS activation in the MPG following chemogenetic manipulation of tumor-responsive DMV neurons. Neuronal somata, shown in the green channel, were identified based on typical ganglionic morphology and intrinsic autofluorescence in the 488-nm channel. Activation of DMV neurons with hM3Dq increased the proportion of FOS⁺ MPG neurons, whereas inhibition with hM4Di reduced MPG neuronal activation. n = 6 mice. **E,F,** Representative fluorescence images (E) and quantification (F) of FOS activation in the MPG following chemogenetic manipulation of tumor-responsive PVN neurons. Activation of PVN neurons with hM3Dq reduced the proportion of FOS⁺ MPG neurons, whereas inhibition with hM4Di increased MPG neuronal activation. n = 5 mice. **G,** Schematic of the experimental design for assessing whether the MPG mediates DMV-driven prostate tumor growth. Ai14/FOS-CreERT2 mice underwent bilateral MPG excision on day 1, followed by injection of AAV-DIO-hM3Dq-EGFP into the DMV on day 7 and orthotopic implantation of RM-1 cells on day 14. CNO and tamoxifen were administered according to the indicated schedule, and tumors were collected on day 38. **H,I,** Representative images of prostate tumors (H) and quantification of tumor weight, tumor volume and body weight (I) in control mice, mice subjected to bilateral MPG excision, and MPG-excised mice with chemogenetic activation of tumor-responsive DMV neurons. MPG excision significantly reduced tumor weight and volume and reduced the tumor-promoting effect of chemogenetic DMV activation. n = 6 mice per group. Data are presented as mean ± SD. One-way ANOVA followed by Tukey’s multiple-comparisons test was used in B, D, F and I. *P < 0.05, **P < 0.01; ***P < 0.001. ns, not significant.

Because the MPG contains activated neurons that innervate prostate tumor and is regulated by the PVN – DMV circuit, we hypothesized that it serves as a key peripheral relay between central autonomic signals and the tumor. Bilateral MPG excision significantly reduced prostate tumor weight and volume (Extended Data Fig. 4I–J). Moreover, after MPG excision, DMV activation did not restore tumor weight or volume to control levels (Fig. 5G–I), indicating that the MPG is a critical downstream mediator of DMV-driven prostate tumor growth.

### WNT3A mediates communication between MPG neurons and prostate tumor

To characterize the transcriptional features of MPG neurons that both innervate prostate tumor and are activated in tumor mice, we combined CTB-EGFP retrograde tracing with FOS-dependent tdTomato labelling in Ai14/FOS-CreERT2 mice. CTB-EGFP⁺/tdTomato⁺ neurons were isolated from the MPG by fluorescence-guided laser microdissection and subjected to Smart-seq2 transcriptomic profiling. We analyzed neurons from normal MPGs (n = 4), tumor MPGs (n = 4) and tumor MPGs with hM3Dq-mediated PVN activation (n = 3). Smart-seq profiles from three groups showed the highest correlations with neurons in a public mouse pelvic ganglion scRNA-seq reference (Spearman ρ = 0.840, 0.651 and 0.748), supporting an overall neuronal transcriptional identity (Extended Data Fig. 5A).

Gene Ontology GSEA showed that, relative to normal MPGs, tumor MPG neurons were depleted for pathways involved in axon fasciculation and transport, synaptic vesicle trafficking, synaptic signalling, neurotransmitter metabolism, postsynaptic organization and neuronal development. By contrast, neuroinflammatory programmes and WNT and STAT signalling were enriched (Fig. 6A). KEGG analysis further revealed enhanced neuroactive ligand–receptor interaction signalling (Fig. 6B-C), indicating tumor-induced transcriptional remodeling characterized by impaired neuronal homeostatic functions, increased inflammation and enhanced intercellular communication.

**Fig. 6.**
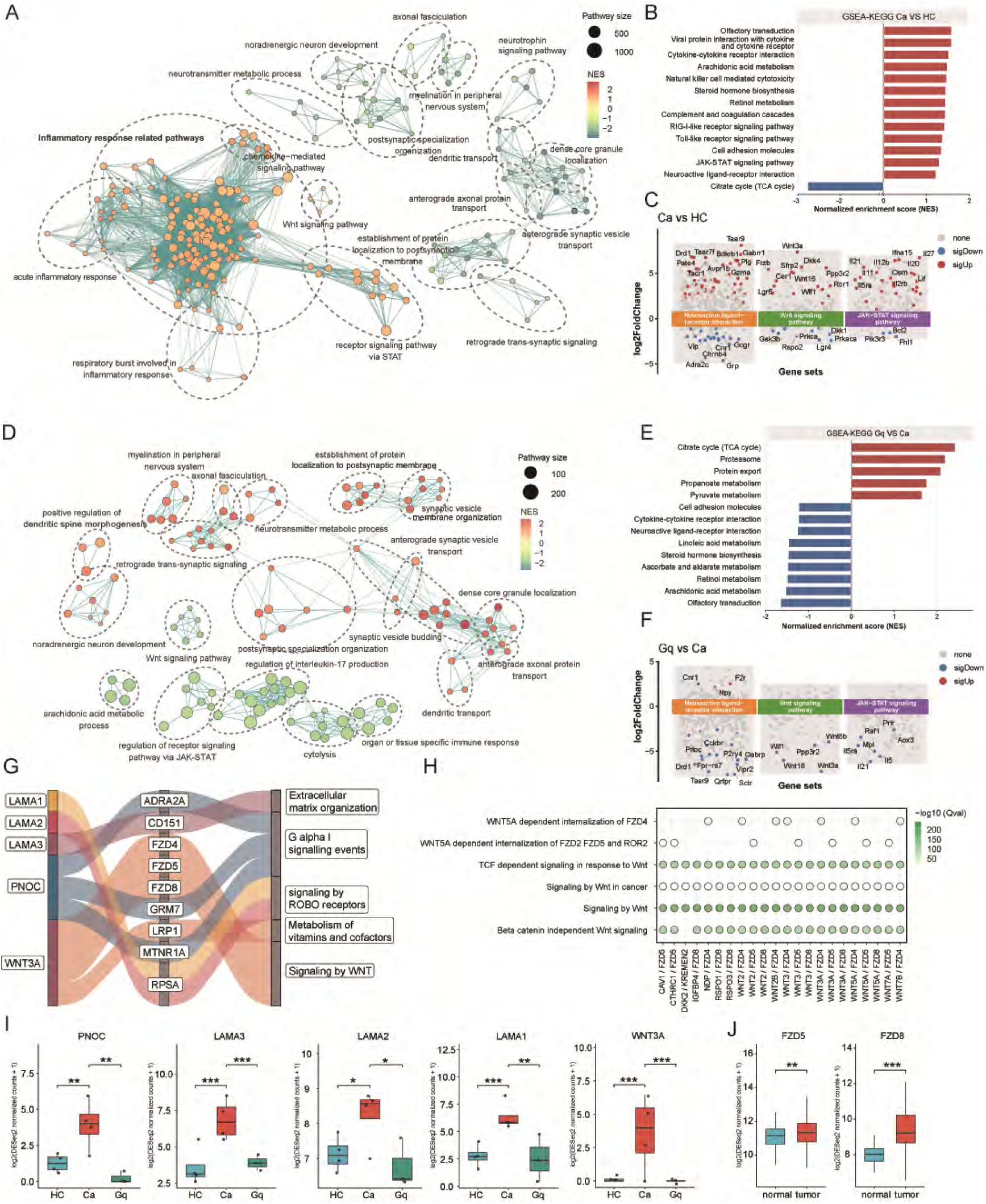
Interaction between MPG neurons and prostate tumor cells via WNT3A. **A–C**, Differential expression and pathway enrichment analyses comparing tumor-innervating, activated MPG neurons from tumor-bearing mice with control. Network plot of significantly enriched GSEA–GO terms. Terms with greater similarity, reflecting a larger number of shared core enrichment genes, are connected by lines, and terms within the same functional cluster are enclosed by dashed outlines (A). Bar plot of significantly enriched GSEA–KEGG pathways (B). Waterfall plots of significant DEGs in the neuroactive ligand–receptor interaction, WNT signalling and STAT signalling pathways (C). **D–F,** Corresponding analyses comparing tumor-innervating, activated MPG neurons following activation of PVN neurons with those from the tumor group. Network plot of significantly enriched GSEA–GO terms (D), bar plot of significantly enriched GSEA–KEGG pathways (E) and waterfall plots of significant DEGs in the indicated pathways (F). **G,** Sankey diagram linking significant ligand–receptor interactions between tumor-innervating, activated MPG neurons and prostate tumor cells to predicted downstream pathway alterations. **H,** Ligand–receptor pairs associated with pathways significantly altered by interactions between MPG neurons and prostate tumor cells. **I,** Box plots comparing ligand gene expression in activated, tumor-innervating MPG neurons among the control, tumor and PVN-manipulated tumor groups. **J,** Box plots comparing WNT3A receptor gene expression between paired prostate tumor and normal samples from the TCGA cohort. DEGs were defined by |log2FC| > 1 and FDR < 0.05. *P < 0.05, **P < 0.01, ***P < 0.001. DEG, differentially expressed gene; FC, fold change; FDR, false discovery rate; GO, Gene Ontology; GSEA, gene set enrichment analysis; KEGG, Kyoto Encyclopedia of Genes and Genomes.

PVN activation mediated by hM3Dq partially reversed this transcriptional programme. GO GSEA showed that, compared with tumor mice, PVN activation enriched neuronal processes including axonal transport, anterograde synaptic vesicle trafficking, neurotransmitter uptake and synaptic vesicle cycling, while inflammatory, WNT and STAT signalling were reduced (Fig. 6D). Neuroactive ligand–receptor interaction signalling was also diminished (Fig. 6E-F), suggesting that PVN activation alleviates tumor-associated remodelling of MPG neurons.

Ligand-receptor analysis predicted interactions between MPG-derived WNT3A and FZD5 or FZD8 expressed by prostate tumor, accompanied by activation of WNT signalling in tumor MPG. These WNT3A-associated interactions were reduced following PVN activation (Fig. 6G–H). Consistently, Wnt3a expression was increased in tumor-associated MPGs and decreased after PVN activation, whereas FZD5 and FZD8 were upregulated in prostate tumor (Fig. 6I–J.) The directions of Wnt3a expression changes remained consistent in leave-one-out sensitivity analysis (Extended Data Fig. 5B). Wnt3a was also rarely detected in public single-cell RNA-sequencing data from normal peripheral ganglia (Extended Data Fig. 5C). Despite overall downregulation of multiple neuronal transport and release modules in the Ca group, Wnt protein secretion pathway genes were significantly upregulated. (Extended Data Fig. 5D, E). Wnt3a ranked highly among extracellular signaling candidates by fold-change magnitude (Extended Data Fig. 5F, G). In contrast, other genes involved in cholinergic, catecholaminergic, and GABAergic transmission showed no significant upregulation in the Ca group (Extended Data Fig. 5H, I).

Together, these findings implicate the WNT3A–FZD axis in MPG neuron–prostate tumor cell communication and support relatively selective tumor-associated Wnt3a induction in MPG neuronal samples.

### Targeting MPG neuronal WNT3A suppresses prostate tumor cell proliferation and migration

Immunofluorescence analysis revealed markedly increased WNT3A signal in activated MPG neurons from tumor mice (Fig. 7A). In normal prostate, WNT3A expression was mainly restricted to the stroma, whereas prostate tumor showed widespread WNT3A expression that either colocalized with or localized adjacent to nerve fibers. FZD8 was also detected near nerve fibers (Extended Data Fig. 6A–B). MPG excision significantly reduced WNT3A protein levels in tumor tissue, whereas DMV chemogenetic activation produced an upward trend (Fig. 7B). Consistently, DMV activation increased the proportion of WNT3A-positive area within the tumor area, whereas DMV inhibition reduced it (Extended Data Fig. 6C). However, MPG excision did not significantly alter tumoral *Wnt3a* mRNA expression (Fig. 7C), whereas in situ hybridization showed increased *Wnt3a* mRNA expression in CHAT^+^ MPG neurons in tumor mice (Fig. 7D), suggesting that the increased WNT3A protein in tumor may partly originate from MPG nerve terminals.

**Fig. 7.**
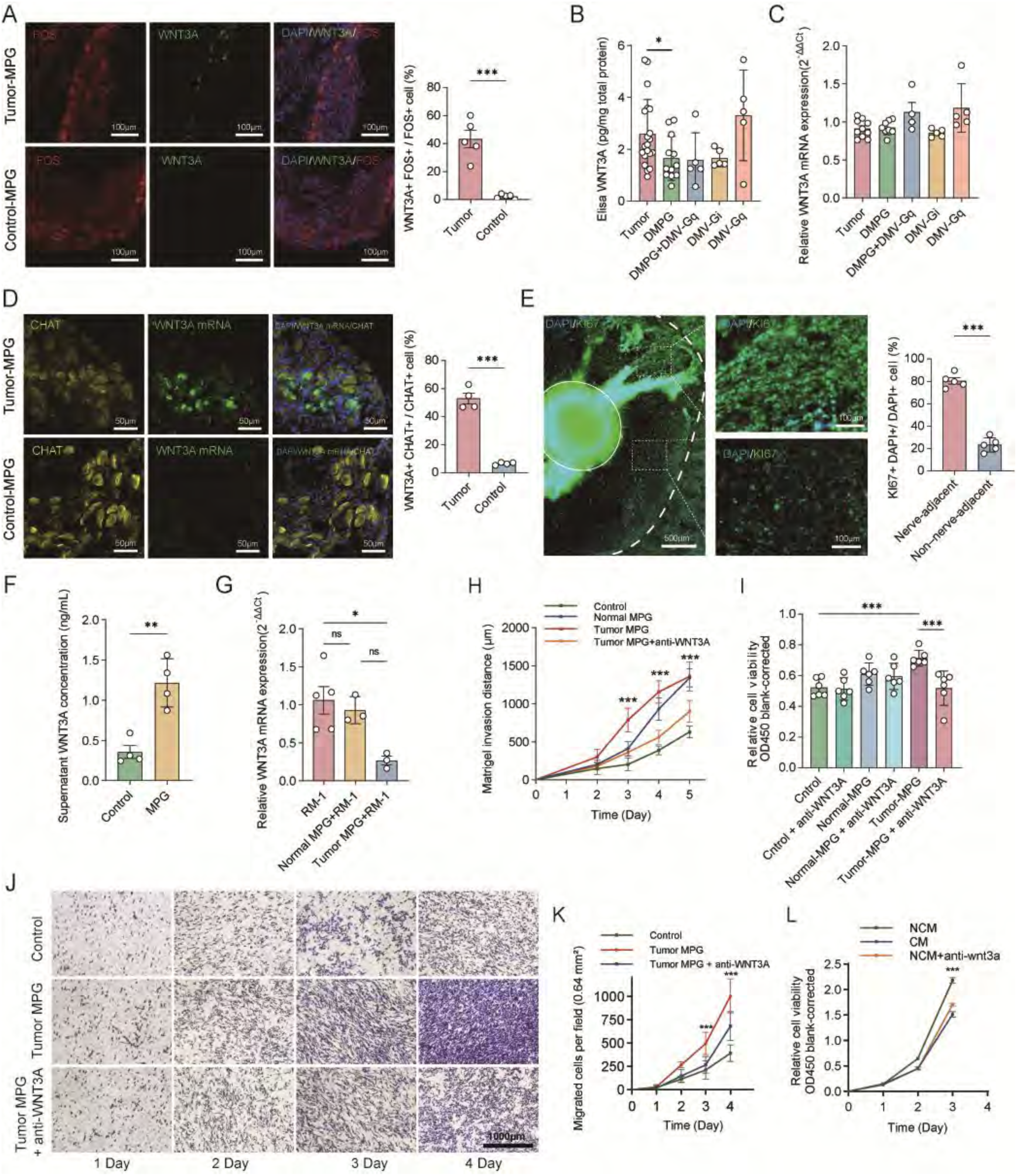
Neutralization of MPG-derived WNT3A suppresses tumor cell proliferation and invasion. **A**, Representative immunofluorescence images and quantification of WNT3A in FOS⁺ MPG neurons from prostate tumor-bearing and healthy control mice. The proportion of WNT3A⁺FOS⁺ cells among FOS⁺ cells was increased in PCa-MPG. n = 5 mice. **B,** ELISA measurement of WNT3A in prostate tumors following MPG excision or chemogenetic manipulation of tumor-responsive DMV neurons. MPG excision reduced WNT3A levels, whereas DMV activation showed an increasing trend. Tumor, n = 20 mice; DMPG, n = 12 mice; DMPG+DMV-Gq, DMV-Gi and DMV-Gq, n = 5 mice per group. **C,** RT–qPCR analysis of Wnt3a expression in prostate tumors following the indicated manipulations. No significant differences were detected among groups. Tumor, n = 10 mice; DMPG, n = 8 mice; DMPG+DMV-Gq, n = 4 mice; DMV-Gi and DMV-Gq, n = 5 mice per group. **D,** Representative fluorescence images and quantification of Wnt3a mRNA in ChAT⁺ MPG neurons. The proportion of Wnt3a⁺ChAT⁺ cells among ChAT⁺ neurons was increased in PCa-MPG. n = 4 mice per group. **E,** Representative fluorescence images of RM-1 cells co-cultured with MPG explants. Tumor cells migrated towards the explants and aligned along nerve fibres, with higher Ki67 expression in nerve-adjacent cells. Dashed and solid circles indicate the Matrigel boundary and MPG explant, respectively. n = 5. **F,** WNT3A concentrations in culture supernatants measured by ELISA. Medium containing MPG explants showed higher WNT3A concentrations than control medium. n = 4 per group. **G,** Relative Wnt3a expression in RM-1 cells cultured alone or with normal or tumor-associated MPG explants. Co-culture with tumor-associated MPG reduced Wnt3a expression in RM-1 cells. n = 3, and 5 for the respective groups. **H,** RM-1 cell invasion distance over time following co-culture with normal or tumor-associated MPG explants, with or without WNT3A-neutralizing antibody. Tumor-associated MPG enhanced invasion, which was attenuated by WNT3A neutralization. n = 6 per group. **I,** Relative viability of RM-1 cells under the indicated co-culture conditions. Tumor-associated MPG increased cell viability, whereas WNT3A neutralization reduced this effect. n = 6 per group. **J, K,** Representative Transwell images (J) and time-course quantification of migrated RM-1 cells (K). Tumor-associated MPG enhanced migration, which was reduced by WNT3A neutralization. n = 8 per group. **L,** Relative viability of RM-1 cells cultured with CM, NCM or NCM supplemented with WNT3A-neutralizing antibody. NCM promoted RM-1 cell proliferation, which was attenuated by WNT3A neutralization. n = 4 per group. NCM, neural conditioned medium derived from MPG explants. Data are presented as mean ± SD. Two-tailed unpaired Student’s t-tests were used in A, D and E. One-way ANOVA followed by Tukey’s multiple-comparisons test was used in B, C, F, G and I. Repeated-measures data in H, K and L were analysed using a linear mixed-effects model, followed by Tukey’s multiple-comparisons test between groups at individual time points. *P < 0.05, **P < 0.01, ***P < 0.001; ns, not significant.

In a co-culture model of tumor cells and MPG explants, tumor cells migrated towards the MPG explants and aligned along nerve fibers, and tumor cells adjacent to nerve fibers exhibited higher Ki67 expression (Fig. 7E and Extended Data Fig. 6D–E). WNT3A levels were significantly higher in supernatants containing MPG explants than in control medium without MPG explants (Fig. 7F). Tumor cells co-cultured with MPG exhibited significantly lower WNT3A mRNA expression than tumor cells cultured alone (Fig. 7G). Co-culture with tumor-associated MPGs increased tumor-cell invasion distance, viability and Transwell migration, all of which were significantly reduced by treatment with WNT3A neutralizing antibody (Fig. 7H–K and Extended Data Fig. 6F).

Isolated glial cells and other MPG-derived adherent outgrowth cells from either normal or tumor-associated MPGs did not significantly enhance tumor cell proliferation (Extended Data Fig. 6G). By contrast, medium containing 25% tumor-associated MPG-conditioned medium promoted tumor cell proliferation, which was significantly suppressed by WNT3A neutralization (Fig. 7L). Together, these findings support a role for MPG-derived WNT3A in promoting prostate tumor cell proliferation and invasion.

### Altered spontaneous brain activity in the human PVN and DMV in prostate cancer

The PVN lies on either side of the third ventricle and participates in neuroendocrine and autonomic regulation. The DMV, located in the dorsal medulla, is a key nucleus for vagal parasympathetic output. Resting-state functional magnetic resonance imaging (rs-fMRI) assesses local spontaneous brain activity through blood oxygen level – dependent signals. Regional homogeneity (ReHo) reflects the temporal synchrony of signals across neighboring voxels, whereas the amplitude of low-frequency fluctuations (ALFF) reflects the magnitude of local low-frequency signal fluctuations.

To investigate whether prostate cancer is accompanied by resting-state functional alterations in these regions, we analyzed rs-fMRI data from 22 patients with prostate cancer (PCa) and 8 patients with benign prostatic hyperplasia (BPH). Age did not differ significantly between the PCa and BPH groups (72.27 ± 7.19 vs 66.75 ± 8.53 years, mean ± SD; t-test, P = 0.131). Voxelwise analysis showed higher indices of local spontaneous brain activity in the PCa group than in the BPH group. ReHo was significantly higher in the PCa group in 12 of the 18 labeled voxels in the PVN and adjacent regions (Extended Data Fig. 7A, B). ALFF was significantly higher in the PCa group in 8 of the 16 labeled voxels in the DMV and adjacent regions (Extended Data Fig. 7C, D).

## Discussion

In this study, we identify an aberrantly activated PVN–DMV–MPG circuit associated with prostate tumor growth. DMV activation enhanced MPG activity, while MPG-derived fibers innervated the tumor microenvironment and supported tumor growth in mice. By combining LMD with Smart-seq2 to profile tumor-innervating and activated MPG neurons, we identified marked transcriptional remodeling, including increased Wnt3a expression. This transcriptional state was modulated by upstream central activity. DMV activation further increased WNT3A in the MPG, and MPG-derived WNT3A promoted prostate cancer cell proliferation and invasion.

Recent advances in cancer neuroscience have shifted the field from a descriptive view of local tumor innervation to a framework in which the brain and peripheral tumors engage in bidirectional and experimentally tractable communication. In peripheral cancers, defined central nodes, including the VTA reward system, ventrolateral medullary catecholaminergic neurons, hypothalamic oxytocin neurons and limbic circuits, regulate tumor growth through autonomic, neuroendocrine and immune relays^16,21–25^. A recent lung adenocarcinoma study further established a bidirectional vagal sensory-to-brainstem-to-sympathetic circuit that restrains antitumor immunity^26^. Our findings extend the brain-tumor circuit framework to a hypothalamic-brainstem-pelvic ganglion axis associated with prostate cancer. Rather than representing only a generalized stress response, this pathway nominates the MPG as a potentially organ-related peripheral relay through which central autonomic state may shape tumor-innervating neurons and prostate tumor progression.

The DMV is a major vagal preganglionic motor nucleus within the dorsal vagal complex and has established roles in gastrointestinal motility, pancreatic regulation, and broader cardiopulmonary and visceral homeostasis^28,29^. In rodents, classical autonomic innervation of the prostate comprises lumbosacral preganglionic inputs conveyed to the MPG through the hypogastric and pelvic nerves, followed by postganglionic projections from the MPG to the prostate^29,30^. Retrograde tracing from the MPG has localized preganglionic neurons mainly to L6–S1 and L1–L2, although this study focused exclusively on the spinal cord and therefore did not assess potential supraspinal regions^31^. In our experiments, focal MPG injection of DiI, a membrane-associated rather than conventional transsynaptic tracer, labelled DMV neurons, supporting a putative DMV-to-MPG projection^32^. Although DiI labelling alone cannot determine whether the labelled axons terminate within the MPG or form synapses with MPG neurons, DMV inhibition and activation respectively decreased and increased MPG neuronal activity, providing functional support for a brainstem-to-pelvic ganglion pathway.

Our findings reveal opposing roles of the PVN and DMV in prostate tumor growth. Although both the PVN and DMV were linked to prostate tumor and showed increased FOS expression during tumor growth, PVN activity restrained, whereas DMV activity promoted tumor growth. This functional antagonism is consistent with broader evidence that the effects of PVN activity vary across cancer models. The PVN is a key hypothalamic node that integrates stress, neuroendocrine and autonomic signals^13,33^. PVN effects on cancer vary across cell types and over time. OXT-expressing PVN neurons suppress colitis-associated colorectal cancer by reducing CG-SMG sympathetic activity^16^. Repeated activation of CRH-expressing PVN neurons promotes lung tumor growth through corticosterone-associated immune suppression, whereas circadian phase-matched activation of the same genetically defined population restores glucocorticoid rhythmicity and restrains breast cancer^17,18^. Tumor-derived LIF and Gal3 also induce a shared brain response involving the PVN and support sympathetic immunosuppression, although that study did not establish PVN-specific causality^15^. PRV tracing has revealed a multisynaptic connection between the PVN and prostate, while PVN transplantation of beta-endorphin neurons attenuated experimental prostate carcinogenesis^34,35^. Our findings extend this framework by implicating the DMV and MPG as downstream components of a PVN-centered circuit regulating prostate tumor growth.

Importantly, concurrent FOS induction in the PVN and DMV does not imply functional equivalence. A plausible interpretation is that the tumor recruits a growth-promoting DMV effect while simultaneously engaging a compensatory PVN response that limits this output. This model is compatible with the diverse effects of the PVN on DMV neurons. Electrical PVN stimulation excites subsets of gut-related DMV neurons through AMPA-mediated glutamatergic transmission but inhibits another subset of DMV neurons through a GRP-sensitive mechanism^36,37^. Conceptually, our findings support a prostate-associated central autonomic checkpoint that places local nerve-tumor interactions within an upstream hypothalamic-brainstem network. Prostate tumors recruit both the PVN and DMV, but these nuclei exert opposing effects, with the PVN acting as a compensatory brake on a pro-tumor DMV-MPG output.

Transcriptomic data for tumor-associated peripheral neurons remain scarce because their somata reside in distant ganglia and are missed by tumor-tissue sequencing. Previous pancreatic and oral cancer studies combined retrograde tracing with neuronal isolation and RNA sequencing but did not enrich neurons activated in vivo^38–40^. Using dual CTB and FOS guided LMD and pooled low-input Smart-seq2, we characterized the transcriptional features of MPG neurons defined simultaneously by anatomical location, prostate-tumor innervation and activity history. Our findings show that the DMV regulates prostate tumor growth through the MPG. DMV activation increased FOS expression in MPG neurons, whereas DMV inhibition reduced it. Bilateral MPG ablation suppressed tumor growth and abolished DMV-induced tumor promotion, establishing the MPG as a necessary downstream mediator. Tumor-innervating, activity-tagged MPG neurons also showed reduced neuronal homeostatic programmes and enrichment of neuroinflammatory and neuroactive ligand–receptor pathways. Consistent with these findings, pancreatic and oral tumors have been shown to reprogramme the sensory or sympathetic neurons that innervate them^38,39^. Other cancer models have revealed neuronal mitochondrial dysfunction, disrupted calcium homeostasis, impaired electrical conduction, axonal or myelin injury, and ATF3–IL-6–type I interferon-associated inflammatory repair programmes^41,42^. Together, these observations and our study support tumor-associated molecular plasticity as an important feature of peripheral neurons.

The transcriptional signatures of impaired homeostasis and inflammation observed in MPG neurons may reflect both retrograde tumor-derived cues and abnormal upstream autonomic input. Previous studies have shown that defined populations of central autonomic neurons can alter neuronal firing or c-FOS expression in downstream autonomic ganglia^26,43^. In a non-cancer stroke model, disruption of the PVN-to-DMV pathway induced α7nAChR–HMGB1-dependent inflammatory changes in pulmonary parasympathetic ganglia^44^. In sensory ganglia, neural activity can be converted into local inflammatory signalling through ATP–P2X7 signalling in satellite glia and neuron–macrophage communication^45,46^. Notably, PVN activation partially shifted the tumor-associated transcriptional programmes towards the healthy state, linking upstream central autonomic activity to the molecular state of MPG neurons. The MPG may therefore represent a downstream node whose activity and transcriptional state are regulated within the PVN–DMV–MPG axis.

MPG remodeling was characterized by tumor associated upregulation of WNT3A expression. Previous studies of prostate cancer neurosignalling have focused primarily on adrenergic and cholinergic pathways, whereas WNT3A has been implicated in AR-related transcription, proliferation, invasion and therapy resistance, but these effects were studied using exogenous WNT3A or in tumor and stromal contexts rather than from a neuronal source^5,47–50^. Previous work provides a physiological basis for neuronal WNT3A expression and Wnt secretion. Hippocampal neuronal systems have been shown to release WNT3A^51,52^. Injury and bone cancer also upregulate WNT3A in DRG neurons^53^. In addition, A-fiber DRG neurons can secrete WNT5A through GPR177^54^. In our study, WNT3A was expressed at low levels in normal MPG neurons but was markedly upregulated in tumor-associated MPG neurons. MPG explants released WNT3A into the culture medium and promoted prostate tumor-cell proliferation and invasion, whereas WNT3A neutralization abolished the proliferative effect of co-culture with MPG explants. Thus, WNT3A was not merely a marker of MPG remodelling; extracellular WNT3A was required for the MPG-associated proliferative effect. Together, these findings support tumor-associated MPG neurons as a previously unrecognized source of WNT3A and identify a functional link between abnormal ganglionic transcriptional states and prostate tumor-cell growth. More broadly, they extend prostate cancer neurosignalling beyond classical neurotransmitters and neuropeptides to a neuron-derived canonical Wnt ligand.

Several limitations should be considered. First, further studies are required to determine whether DMV axons terminate within the MPG and form direct synapses with MPG neurons. Additional circuit routes outside the proposed PVN – DMV – MPG axis also cannot be excluded. Second, WNT3A detected in conditioned medium from MPG explants supports ganglion-level release ex vivo, but its cellular source and release from tumor-innervating terminals remain unresolved. Third, pooled Smart-seq2 enabled targeted profiling of tumor-innervating, activity-tagged MPG neurons but provides limited resolution of cellular heterogeneity and rare transcripts. Fourth, region-level manipulation of tumor-activated neurons in the PVN or DMV altered tumor growth but cannot exclude contributions from parallel autonomic, endocrine or immune pathways. Finally, validation in additional prostate cancer models and human tissues will be required, given the differences between the RM-1 model, rodent MPG anatomy and human disease.

Together, our findings identify a polysynaptic PVN–DMV–MPG circuit that regulates prostate tumor growth and implicate WNT3A as a mediator of the pro-tumor effects of the MPG (Extended Data Fig. 8), highlighting the therapeutic potential of targeting the neural circuit and downstream molecular pathway.

## Methods

### Clinical specimens

Prostate tumour tissue samples for single-cell RNA sequencing were obtained from six male patients recruited at Shanghai East Hospital Affiliated to Tongji University between July 2023 and March 2024. Patients were eligible if they were aged 40–90 years, had biopsy-confirmed metastatic prostate cancer, and had received no previous anticancer or surgical treatment.

The MRI analysis included 22 male patients with prostate cancer selected from an ongoing prospective observational cohort registered at ClinicalTrials.gov (NCT05737004), together with 8 male patients with benign prostatic hyperplasia (BPH) who served as controls. Participants in the MRI analysis were prospectively recruited at the same institution between March 2023 and September 2026. For the present analysis, patients with prostate cancer were eligible if they were aged 18–80 years, had newly diagnosed localized prostate cancer, and had received no previous anticancer or surgical treatment. Patients with BPH were eligible if they were aged 18–80 years and had a clinical diagnosis of BPH. Exclusion criteria for both groups included brain atrophy, cerebral ischemia, traumatic brain injury, epilepsy, brain tumours, previous brain surgery, or psychiatric disorders. This exploratory MRI analysis was not prespecified in the original study protocol or statistical analysis plan and was undertaken to explore the clinical relevance of the experimental findings. Recruitment to the registered cohort was ongoing at the time of this analysis.

All participants provided written informed consent before participation. The single-cell RNA-sequencing study and the MRI study were approved by the Medical Ethics Committee of Shanghai East Hospital (approval ID: [2022] Research Review No. 121 and [2020] Research Review No. 082).

### Single-cell RNA sequencing and data processing of clinical specimens

Tumor tissues were processed into single-cell suspensions, and 3′ single-cell transcriptome libraries were generated using the 10x Genomics Chromium platform. Libraries were sequenced as 150-bp paired-end reads on an Illumina NovaSeq 6000 instrument by Shanghai NEO-Bio Biotechnology Co., Ltd. Raw sequencing data were processed using Cell Ranger v3.0 and subsequently imported into Seurat for downstream analysis. Cells with 200–7,500 detected genes and a mitochondrial gene fraction below 20% were retained, and doublets were removed using DoubletFinder. Following normalisation, highly variable gene selection and principal-component analysis (PCA), Harmony was used for batch correction. Cells were then clustered and visualised using uniform manifold approximation and projection (UMAP). Tumor and normal epithelial cell populations were identified and retained for downstream analyses.

Within each tumor-cell or normal epithelial-cell subpopulation, differential expression between the two groups was assessed using the Wilcoxon rank-sum test. Genes with an adjusted P value below 0.05 and an absolute average log₂ fold change greater than 0.25 were considered differentially expressed. Gene Ontology (GO) Biological Process enrichment analysis was performed using clusterProfiler. Module scores for selected gene sets were calculated using the Seurat AddModuleScore function to provide an integrated measure of the normalised expression of the genes within each gene set. Based on previous studies, three neural-related gene signatures were used: a neuronal cell marker signature, a perineural invasion (PNI) marker signature, and a composite C-Neural signature^55^. The C-Neural signature integrated neural-related genes from KEGG, GO, and literature-derived neuronal cell and PNI markers. Gene sources and selection criteria are provided in Extended Data Table 1, and the complete gene lists are provided in Extended Data Table 2.

Transcriptomic and clinical data from 493 patients with prostate cancer were obtained from The Cancer Genome Atlas (TCGA) through the National Cancer Institute Genomic Data Commons (GDC) Data Portal^56^. The first biochemical recurrence (BCR) was used as the study endpoint, and days to first biochemical recurrence was used as the time-to-event variable. For each gene, the optimal expression cutoff was determined using maximally selected rank statistics, and patients were accordingly classified into high- and low-expression groups. Kaplan–Meier analyses were then performed to evaluate BCR-free survival.

### MRI Protocol

MRI data were acquired using a 3.0 T MR (MR750w, GE Healthcare, United States). Scanning was performed by Yibin Wang, who had >10 years of professional experience. Participants kept their eyes closed during the resting-state acquisition. Resting-state functional images (Rs-fMRI) were acquired using a T2*-weighted gradient-echo echo-planar imaging sequence with the following parameters: repetition time (TR), 2000 ms; echo time (TE), 30 ms; flip angle, 90°; field of view (FOV), 220 × 220 mm²; acquisition matrix, 64 × 64; slice thickness, 3 mm; interslice gap, 1 mm; and 36 axial slices per volume. A total of 180 volumes were acquired per participant. Three-dimensional T1-weighted structural images were acquired using an inversion-prepared gradient-echo sequence with the following parameters: TR, 8 ms; TE, 3 ms; inversion time, 1100 ms; flip angle, 7°; acquisition matrix, 256 × 256; nominal slice thickness, 1 mm; and FOV, 240 × 240 mm². Images were reconstructed with a 512 × 512 matrix and approximately 0.5-mm spacing between reconstructed slices.

### Rs-fMRI Preprocessing

Resting-state fMRI data were preprocessed using the advanced edition of the Data Processing Assistant for Resting-State fMRI (DPARSFA, version 5.2), within DPABI^57^. Image processing was performed by LW, who had 4 years of experience. DICOM images were converted to NIfTI format, and the first 10 volumes were discarded, leaving 170 volumes per participant. Slice-timing correction was performed for 36 slices acquired in an interleaved order, using the 35th slice as the reference. Head motion was corrected using six-parameter rigid-body realignment with a two-pass registration-to-mean procedure. T1-weighted images were co-registered to the mean functional image and segmented into gray matter, white matter, and cerebrospinal fluid. Functional images were normalized to Montreal Neurological Institute space using DARTEL and resampled to 3 × 3 × 3 mm³ voxels. Subsequent processing was performed separately for ALFF and ReHo. Both branches included linear detrending and nuisance regression using the Friston-24 motion model and mean white-matter and cerebrospinal-fluid signals extracted from standard tissue masks supplied with DPABI and resampled to the functional-image grid. Global signal regression and motion scrubbing were not performed.

For ALFF analysis^58^, the normalized functional images were spatially smoothed using a Gaussian kernel with a full width at half maximum of 4 mm before linear detrending and nuisance regression. The voxelwise time series were transformed into the frequency domain using a fast Fourier transform. ALFF was calculated as the mean amplitude within the 0.01–0.10 Hz frequency band. ALFF estimation was performed before temporal band-pass filtering, and individual ALFF maps were retained for subsequent voxelwise statistical analysis. For ReHo analysis^59^, the normalized functional images were processed without prior spatial smoothing. Following linear detrending and nuisance regression, the time series were band-pass filtered at 0.01–0.10 Hz. ReHo was calculated using Kendall’s coefficient of concordance for the time series within a 27-voxel neighborhood comprising each voxel and its 26 adjacent voxels, restricted to the analysis brain mask. Both ReHo and ALFF maps were spatially smoothed after computation and subsequently used for voxelwise statistical analyses.

The bilateral hypothalamic paraventricular nucleus (PVN) mask was derived by combining labels 21 and 22 from the 0.5-mm isotropic hypothalamic atlas of Neudorfer et al.^60^, distributed in MNI152 NLIN 2009b space via Zenodo (version 4). The region encompassing the dorsal motor nucleus of the vagus (DMV) was defined using the viscero-sensory-motor (VSM) nuclei complex atlas of Singh et al.^61^, obtained from Brainstem Navigator v1.0. Based on this atlas, the DMV was manually outlined for anatomical illustration with reference to previous studies^62^. Voxelwise between-group differences in ReHo and ALFF within these masks and their neighboring voxels were assessed using ordinary least-squares regression, with diagnostic group and mean-centered age as predictors. Group effects were tested using heteroscedasticity-consistent HC3 standard errors, with two-sided P values calculated from a t-distribution with n−3 degrees of freedom. Results were evaluated at an uncorrected threshold of P < 0.05. Image preprocessing and calculation of ReHo and ALFF were performed using DPARSFA and SPM12 in MATLAB. NIfTI images and ROI masks were processed using NiBabel, and anatomical overlays and voxelwise statistical maps were generated using Matplotlib, both within the Python environment.

### Mice

Male mice were used throughout the study. Specific-pathogen-free male C57BL/6J mice (8 weeks old) were purchased from Beijing Vital River Laboratory Animal Technology Co., Ltd. (Beijing, China). Ai14/FOS-CreERT2 mice (8-10 weeks old) were kindly provided by Prof. Gang Chen at Zhejiang University School of Medicine. Mice were housed under controlled conditions at 24 ± 1 °C and 50–70% relative humidity, with a 12-h light/dark cycle and lights on from 07:00 to 19:00. All animal procedures complied with the Regulations for the Administration of Affairs Concerning Experimental Animals of China and were approved by the Animal Ethics and Welfare Committee of Shanghai East Hospital, Tongji University (Approval ID: TJBB08126101).

### General surgical procedures

Mice were anaesthetised with 3% isoflurane in an induction chamber, and anaesthesia was maintained with 1.5–2% isoflurane delivered through a nose cone or via endotracheal intubation connected to a ventilator. Animals were placed on a heating pad, and their body temperature was maintained at approximately 36–37 °C throughout the procedure. Ophthalmic ointment was applied to prevent corneal drying. Following surgery, mice received supplemental oxygen on the heating pad until they regained consciousness.

### Orthotopic prostate tumor model

RM-1 prostate cancer cells (iCell-0105a, Cellverse) were cultured in DMEM supplemented with 10% fetal bovine serum (FBS) and 1% penicillin–streptomycin (Gibco) in a humidified incubator at 37°C with 5% CO₂. For orthotopic implantation, RM-1 cells in the logarithmic growth phase were washed with sterile PBS and detached using trypsin–EDTA. After neutralization with complete medium, cells were collected by centrifugation and washed again with sterile phosphate-buffered saline (PBS) to remove residual serum and trypsin. The cells were then resuspended in cold PBS at the required concentration.

Unless otherwise indicated, 8-week-old male C57BL/6J and Ai14/FOS-CreERT2 mice were used. Under a stereomicroscope, the abdominal hair was removed and an approximately 1-cm midline incision was made. The anterior prostate lobe was carefully exposed, and 5 × 10⁴ RM-1 cells were injected into the prostate using a Hamilton microsyringe. The formation of a translucent bleb at the injection site indicated successful tumor-cell implantation. Mice were then returned to their home cages and maintained for a further 2–3 weeks. Mice were humanely euthanized if they exhibited predefined humane endpoints, including >20% body-weight loss, severe lethargy, impaired mobility, abdominal distension, urinary or defecatory difficulty, or a palpable tumor diameter >15 mm. Euthanasia was performed by intraperitoneal overdose of sodium pentobarbital. The resulting prostate tumors were subsequently excised and weighed, and their dimensions were measured using calipers. Tumor volume was calculated as follows: volume = 0.5 × longest diameter × perpendicular diameter².

### Benign prostatic hyperplasia model

To establish the BPH model, Ai14/FOS-CreERT2 mice (6–8 weeks old) underwent castration under anesthesia. After a 7-day recovery period, mice received daily subcutaneous injections of testosterone propionate (5 mg/kg) for 4 consecutive weeks. Three days after cessation of testosterone administration, mice received a single injection of tamoxifen, and tissues were collected 2 days later.

### Activity-dependent neuronal labelling

Ai14/FOS-CreERT2 transgenic mice were used to genetically label activated neurons associated with prostate tumor. After prostate tumor had formed, tumor-associated stimuli induced FOS expression in a subset of activated neurons, thereby driving CreERT2 expression. Tamoxifen (HY-13757A, MedChemExpress) was subsequently administered to induce the nuclear translocation of CreERT2 and recombination at the Ai14 reporter locus. Consequently, FOS-positive neurons activated during the labelling window permanently expressed the red fluorescent protein tdTomato. Tamoxifen dissolved in corn oil was administered intraperitoneally at 50 mg/kg. For experiments examining activated neurons, MPG, IMG, DRG and brain tissues were collected 2 d after tamoxifen administration, fixed and sectioned. tdTomato-positive neurons were visualized and recorded by fluorescence microscopy. For experiments involving brain-region manipulation after tumor implantation, tamoxifen administration began on day 3 after tumor implantation and was continued once weekly for 2–3 weeks, depending on the experimental design.

### Retrograde neuronal labelling

Tumor-bearing mice and normal 8-week-old mice were anaesthetised with isoflurane. The hair overlying the target organ was removed, and an approximately 1-cm incision was made directly above the prostate or tumor.

Following microsurgical exposure, DiI (HY-D0083, MedChemExpress; 0.5%, 10 µL), cholera toxin B subunit–EGFP (CTB–EGFP, BrainVTA; 0.1%; 8 µL) or pseudorabies virus (PRV, BrainVTA; 2.0 × 10⁹ PFU/mL, 150 or 300 nL) was injected slowly into multiple sites in the anterior prostate lobe or prostate tumor using a Hamilton microsyringe. Before wound closure, the injection site was rinsed with 0.5 ml sterile saline to remove any leaked tracer. Mice were euthanised 3–28 d after tracer injection, according to the experimental design, and tissues were collected for immunofluorescence or laser-capture microdissection followed by Smart-seq2 analysis.

To complement the retrograde tracing experiments, we examined anterograde tracing data from the Allen Mouse Brain Connectivity Atlas (experiment ID: 299759881) to assess PVN – DMV projections. This dataset was generated by Cre-dependent AAV-EGFP injection targeting the PVN in an Efr3a-Cre_NO108 mouse. Coronal images were qualitatively inspected using the Allen online viewer to assess EGFP-labelled axonal fibres in the DMV, with anatomical regions identified using the Allen Reference Atlas.

### Dissection of the major pelvic ganglia

After euthanasia, the abdominal cavity was exposed through a lower midline incision. The bladder and seminal vesicles were gently displaced to expose the prostate. Under an illuminated stereomicroscope, the bilateral MPGs were identified adjacent to the dorsolateral prostate. Each ganglion was dissected intact using fine forceps and microscissors and immediately transferred to ice-cold artificial cerebrospinal fluid (ACSF). Intact ganglia were processed for whole-mount staining to quantify neuronal cell types or to assess retrograde labelling.

For direct intraganglionic labelling, glass micropipettes connected to a microinjection pump (L0107-2A; Longer) were used to inject 500 nl DiI (HY-D0083, MedChemExpress; 0.5%) bilaterally into the MPGs at a rate of 200 nl min⁻¹, and the pipette was left in place for several minutes before withdrawal to minimize leakage. For the MPG excision experiment, under anesthesia, the MPGs were exposed and identified on both sides by visualizing their major branches and accompanying vessels. Each MPG was carefully dissected from the surrounding connective tissue and excised while minimizing vascular injury. In the sham-operated group, all surgical procedures before MPG removal were performed in the same manner, but the ganglia were left intact. To validate successful denervation, the excised MPGs were fixed, embedded in paraffin, sectioned, and histologically examined to confirm complete removal.

### Dissection of dorsal root ganglia

After euthanasia, a longitudinal midline incision was made along the back to expose the vertebral column. Under a stereomicroscope, the laminae were removed, the vertebral canal was opened and the spinal cord was exposed. The dorsal roots were followed laterally to identify the DRGs within the intervertebral foramina, and the target DRGs were identified according to their spinal levels. DRGs from T12 to S3 were carefully isolated using fine forceps and microscissors and collected for subsequent immunofluorescence analysis.

### Dissection of the inferior mesenteric ganglion

After euthanasia, the abdominal cavity was opened through a midline incision, and the posterior abdominal wall and abdominal aorta were exposed. Under an illuminated stereomicroscope, the inferior mesenteric artery was first identified at its origin from the abdominal aorta, and the inferior mesenteric ganglion was located near the arterial origin. The ganglion was isolated using fine forceps and microscissors and transferred to ice-cold ACSF for subsequent experiments.

### Stereotactic viral injection into the PVN and DMV

Mice were prepared for surgery as described above. The scalp was disinfected with 75% ethanol, and the skull was exposed. A burr hole approximately 1 mm in diameter was drilled above the P VN at the following coordinates: anteroposterior (AP), −0.93 mm; mediolateral (ML), ±0.25 mm; and dorsoventral (DV), −4.75 mm. A second burr hole was drilled above the DMV at AP −7.32 mm, ML ±0.25 mm and DV −4.40 mm. Viral vectors were delivered through glass micropipettes connected to a microinjection pump (L0107-2A; Longer). At each injection site, 0.2 µl virus was infused at a rate of 0.02 µl min⁻¹. After injection, the micropipette was left in place for 10 min to facilitate viral diffusion and minimise reflux. The micropipette was then withdrawn slowly, and the scalp was sutured. The following viruses were utilized in the experiments: rAAV-EF1a-flex-taCasp3, PRV-EGFP, AAV-hSyn-DIO-hM4Di-EGFP, AAV-hSyn-DIO-hM3Dq-EGFP, AAV-hSyn-DIO-EGFP, AAV-EF1a-flex-EGFP. All viruses were purchased from BrainVTA in Wuhan, China.

Chemogenetic experiments were initiated after sufficient DREADD expression had been established. Clozapine N-oxide (CNO; SD2461, Beyotime) was administered intraperitoneally at 3 mg/kg every 2 d during the experimental period to activate hM3Dq-expressing neurons or inhibit hM4Di-expressing neurons.

### Immunofluorescence

Mice were transcardially perfused through the left ventricle with PBS followed by 4% paraformaldehyde (PFA). Brain and tumor tissues were post-fixed in 4% PFA for 24 h and cryoprotected in 30% sucrose for 72 h. Ganglia were post-fixed in 4% PFA for 3 h and cryoprotected in 30% sucrose overnight. Brain tissues were cryosectioned at 20 µm, whereas ganglia and tumor tissues were sectioned at 10 µm. Cultured neurons were fixed with 4% PFA for 10 min. Sections were permeabilised and blocked for 1.5–2 h in a solution containing 1% Triton X-100 and 5% bovine serum albumin and incubated with primary antibodies for 24 h at 4 °C. The following primary antibodies were used: rabbit anti-OXT (1:400; ab212193, Abcam), rabbit anti-ChAT (1:400; GB150271, Servicebio), mouse anti-TH (1:400; GB15181, Servicebio), mouse anti-TUBB3/TUJ1 (1:400; 66375-1-Ig, Proteintech), rabbit anti-WNT3A (1:500; ab219412, Abcam), rabbit anti-FZD8 (1:400; BS2700, Bioworld), rabbit anti-OXTR (1:400; GB112522, Servicebio), rabbit anti-AChE (1:400; GB15038, Servicebio) and rabbit anti-Ki-67 (1:200; 84432-1-RR, Proteintech). Sections were incubated with Alexa Fluor 488-, 594- or 647-conjugated secondary antibodies for 1 h at room temperature and mounted using an antifade mounting medium containing 4 ′, 6-diamidino-2-phenylindole (DAPI; Beyotime Biotechnology). Images were acquired using a Leica fluorescence microscope and analysed using ImageJ (1.53t; National Institutes of Health, Bethesda, MD, USA).

For each sample, two to three non-adjacent sections containing the anatomically matched target brain region were selected according to the mouse brain atlas. For each ganglion, serial sections were screened to identify the section with the largest cross-sectional ganglion profile. This reference section, together with two non-adjacent flanking sections along the serial sectioning axis, was selected for analysis. The entire region of interest was outlined, and all marker-positive neuronal somata within the ROI were counted by an investigator blinded to group allocation. Values from sections of the same sample were averaged for statistical analysis.

### SweAMI RNA fluorescence in situ hybridisation

After collection, MPGs were rinsed in PBS and immediately fixed in in situ hybridisation fixative for at least 12 h at 4 °C. Tissues were then sequentially dehydrated in 15% sucrose for 8 h and 30% sucrose overnight. Cryosections were air-dried at room temperature and re-fixed in in situ hybridisation fixative for 10 min. Following PBS washes, the sections underwent tissue retrieval and proteinase K digestion. Prehybridisation solution was then applied for 1 h at 40 °C, followed by overnight hybridisation with target-gene-specific probes under controlled-temperature conditions.

After hybridisation, the sections were washed using a graded series of saline–sodium citrate (SSC) solutions. Branched probes were sequentially hybridised for 45 min at 40 ° C, followed by incubation with fluorescent signal probes (1:200) for 3 h at 40 °C to amplify the SweAMI signal. After SSC washes, nuclei were counterstained with DAPI for 8 min, and the sections were mounted using antifade mounting medium. Fluorescence signals were visualised and imaged using a Nikon upright fluorescence microscope.

Three gene-specific oligonucleotide probes targeting mouse Wnt3a mRNA (NCBI Gene ID: 22416) were designed for SweAMI fluorescence in situ hybridization: probe 1, 5 ′ - ACTAAGAGGTATCCGAGAGGAGCC-3′; probe 2, 5′-ATGGCATGGACAAAGGCTGACTCC-3′; and probe 3, 5′-CTCCCGAGAGACCATTCCTCCAAA-3′.

### Fluorescence-guided laser-capture microdissection and Smart-seq2 profiling of targeted neurons

To isolate activated MPG neurons that innervated prostate tumor, CTB–EGFP was injected into the prostate tumor of Ai14/FOS-CreERT2 mice to retrogradely label tumor-innervating neurons. Tamoxifen was administered intraperitoneally to induce FOS-CreERT2-mediated tdTomato expression and thereby label FOS-activated neurons. Intact MPGs were harvested and cut into 10-μm-thick cryosections. Under fluorescence guidance, CTB–EGFP- and tdTomato-double-labeled neurons were identified and excised using a laser microdissection system. For each sample, all neurons within the entire MPG that met these fluorescence criteria were dissected and pooled as an independent biological sample for subsequent transcriptomic analysis.

Total RNA was extracted from the laser-microdissected neurons. Smart-seq2 was used for low-input transcriptome amplification and library construction. Briefly, RNA was reverse-transcribed using oligo(dT) primers, and template switching was mediated by a locked-nucleic-acid-containing template-switching oligonucleotide to improve the capture of low-abundance transcripts and the efficiency of cDNA amplification. The resulting cDNA was preamplified, purified and subjected to quality control before sequencing-library construction. Libraries were sequenced as 150-bp paired-end reads on an Illumina NovaSeq platform.

Raw sequencing data were quality controlled using fastp v0.20.0 to generate clean reads. Reads were aligned to the mouse reference genome using HISAT2 v2.2.1, and alignment files were sorted using SAMtools v1.15.1. Gene-level raw read counts were generated using featureCounts v2.0.1.

Gene-level raw counts from Smart-seq2 were analysed using DESeq2. The analysis included MPG neuronal samples from healthy control mice (n = 4), prostate tumour-bearing mice (n = 4), and tumour-bearing mice in which tumour-responsive PVN neurons were chemogenetically activated using hM3Dq (n = 3). All pairwise comparisons among the three groups were performed. Genes with a Benjamini – Hochberg-adjusted P value (false-discovery rate, FDR) below 0.05 and an absolute log₂ fold change greater than 1 were considered significantly differentially expressed.

Upregulated and downregulated genes were separately subjected to over-representation analysis using clusterProfiler for GO Biological Process, Cellular Component and Molecular Function terms and mouse Kyoto Encyclopedia of Genes and Genomes (KEGG) pathways. In addition, GO and KEGG gene-set enrichment analysis (GSEA) was performed using all genes ranked in descending order of log₂ fold change. aPEAR was used to cluster neuronal-function-related GO Biological Process pathways according to their similarity and to visualise the resulting enrichment networks.

Focused analyses were further performed for the KEGG neuroactive ligand–receptor interaction pathway (mmu04080) and the GO Biological Process pathways Wnt signaling pathway (GO:0016055) and JAK−STAT signaling pathway (GO:0007259). Within these pathways, genes with an FDR below 0.05 and an absolute log₂ fold change greater than 0.5 were selected and visualized using waterfall plots.

### Ligand – receptor interaction network construction and key signalling-pathway screening based on Smart-seq2 data

To evaluate potential communication between prostate tumor and MPG neurons, BulkSignalR was used to integrate mouse MPG Smart-seq2 data with human prostate transcriptomic data from TCGA–PRAD. Mouse genes were first converted to their human orthologues, and the MPG and prostate datasets were integrated on the basis of shared genes. Before communication inference, differential expression analysis was performed using DESeq2, comparing tumor-bearing MPG samples with healthy MPG controls and TCGA–PRAD tumor samples (n = 497) with normal prostate samples (n = 52). The resulting differential expression statistics were used as input for BulkSignalR.

Communication was inferred in the MPG-to-prostate-tumor direction. BulkSignalR integrates known ligand–receptor interactions from LRdb with GO Biological Process and Reactome pathway information to construct ligand–receptor–pathway axes comprising ligands, receptors and receptor-associated downstream target genes. Statistical support was assessed according to the coordinated changes in these three components under the corresponding disease conditions, using an FDR q value of 0.10. For each ligand–receptor pair, the pathway with the strongest support was retained. Communication-axis activity scores were subsequently calculated, and functional enrichment analysis was performed for the associated target genes.

### Co-culture of MPG explants and tumor cells

Freshly isolated MPGs were immediately embedded in 8-µl Matrigel drops in 12-well plates. After incubation at 37 °C for 10 min to allow the Matrigel to polymerise, DMEM supplemented with 10% fetal bovine serum was added slowly to each well. MPGs were maintained at 37 °C in 5% CO₂ for 48–72 h until prominent neurite outgrowth was observed.

Exponentially growing tumor cells were resuspended at a concentration of 1 × 10⁷ cells ml⁻¹, and 8 µl of the cell suspension was added around each MPG-containing Matrigel drop. Plates were incubated at 37 °C for 10 min to allow cell attachment, after which complete medium was added slowly along the wall of each well. Co-cultures were maintained for 24–72 h. Tumor-cell infiltration into the Matrigel region and MPG neurite outgrowth were observed and imaged by microscopy.

MPG-derived glial cells were obtained using an explant-outgrowth method. Freshly dissected MPGs were embedded in small droplets of Matrigel. After polymerization at 37 °C, complete culture medium was carefully added, and the explants were maintained at 37 °C in a humidified atmosphere containing 5% CO₂. The medium was replaced every 2 d. After the cells migrating from the MPG explants reached 70–80% confluence, the explants and residual Matrigel were removed. The adherent outgrowth cells were detached using 0.25% trypsin–EDTA and replated for subsequent experiments. The medium was replaced 1 d after replating and every 2 d thereafter.

### Transwell assay of MPG explant and tumor cells

Freshly isolated MPGs were immediately embedded in 8 µl Matrigel in the lower chambers of 12-well Transwell plates (FTW043, Beyotime). After incubation at 37 °C for 10 min to allow the Matrigel to polymerise, the explants were cultured in DMEM supplemented with 10% FBS for 48–72 h. Tumor cells were resuspended in serum-free DMEM at 3 × 10⁵ cells ml⁻¹, and 500 µl of the cell suspension was added to each upper insert. The plates were left undisturbed for 15 min to minimise aggregation while the cells settled.

After co-culture for the indicated periods, the Transwell membranes were washed with PBS, fixed with 4% PFA for 10 min and stained with crystal violet (C0121, Beyotime) for 10 min. Cells remaining on the upper surface of the membrane were gently removed using a cotton swab. Five randomly selected fields were imaged under an inverted microscope, and cells on the lower surface of the membrane were quantified using ImageJ.

### Preparation of MPG-conditioned medium

To collect MPG-conditioned medium (MPG-CM), freshly isolated MPGs were embedded in Matrigel in 12-well plates and cultured for 5 d in DMEM supplemented with 10% FBS and 1% penicillin–streptomycin (Gibco) to establish an axonal network. The medium was replaced on day 5, after which MPG-CM was collected every 2 d for a further 6 d. The collected medium was passed through a 0.22-µm membrane filter and stored at −80 °C until use.

### WNT3A neutralisation

For WNT3A-neutralisation experiments, Wnt-3a-neutralising antibody (MAB9025; R&D Systems) or a matched rat IgG1 isotype control (MAB005; R&D Systems) was added to the conditioned medium at the time of tumor-cell seeding to a final concentration of 1 µg ml⁻¹. The medium was replaced daily, and the respective antibodies were replenished at each medium change to maintain the same concentration.

### CCK-8 assay

RM-1 cells were seeded in 96-well plates at 1,500 cells per well and cultured in medium containing 25% MPG-CM. After 24, 48, or 72 h, the culture medium was replaced with 100 µL serum-free medium containing 10 µL Cell Counting Kit-8 reagent (CCK-8; ATYH12131, Abbkine) per well. The plates were incubated at 37 °C for 1 h, after which absorbance at 450 nm (OD450) was measured using a microplate reader. Wells containing the same medium and CCK-8 reagent but no cells were used as blanks for background correction.

For co-culture experiments, tumor cells were dissociated and collected after 3 d of co-culture with MPGs and seeded in 96-well plates at 20,000 cells per well. After allowing the cells to attach for 3 h, the medium was removed, and the CCK-8 assay was performed as described above.

### Enzyme-linked immunosorbent assay

Fresh mouse tumor tissues were pulverised under liquid nitrogen and weighed. The tissue powder was homogenised in nine volumes of PBS using a tissue homogeniser. The supernatant was collected for subsequent analysis.

WNT3A concentrations in culture supernatants and mouse prostate tumor were measured using an enzyme-linked immunosorbent assay kit (EM0737; FineTest) according to the manufacturer’s instructions. Briefly, samples and standards were added to antibody-coated plates. After incubation and washing, the enzyme-labelled detection antibody, horseradish peroxidase–streptavidin complex and tetramethylbenzidine substrate solution were added sequentially. The optical density of each well was measured at 450 nm using a microplate reader (PerkinElmer), and WNT3A concentrations were determined from the resulting standard curve.

### RNA extraction and quantitative real-time PCR

Total RNA was extracted from mouse tumor tissues and cultured tumor cells using RNA Extraction Solution (G3013; Wuhan Servicebio Technology Co., Ltd., China). A total of 2 µg RNA was reverse-transcribed into cDNA using SweScript All-in-One RT SuperMix for qPCR (G3337; Servicebio). Quantitative real-time PCR was performed using 2× Universal Blue SYBR Green qPCR Master Mix (G3326; Servicebio) on a QuantStudio 5 Real-Time PCR System (Applied Biosystems, USA). Gapdh was used as the reference gene, and relative Wnt3a expression was calculated using the 2^−ΔΔCt^ method.

The primer sequences were as follows: mouse Wnt3a (NM_009522.2), forward 5 ′ - ATCTGGTGGTCCTTGGCTGTG-3′ and reverse 5′-ACTCCTGGATGCCCGCTTT-3′, yielding a 166-bp product; and mouse Gapdh (NM_008084.2), forward 5 ′ - CCTCGTCCCGTAGACAAAATG-3 ′ and reverse 5 ′-TGAGGTCAATGAAGGGGTCGT-3 ′, yielding a 133-bp product.

### Statistics

All statistical analyses were performed using R version 4.5.3 (R Foundation for Statistical Computing, Vienna, Austria). For univariate analyses, comparisons between two groups were performed using two-tailed unpaired Student’s t-tests, whereas comparisons involving three or more groups were performed using one-way ANOVA followed by Tukey’s multiple-comparisons test. Repeated-measures data were analysed using linear mixed-effects models, followed by Tukey’s multiple-comparisons tests between groups at individual time points. Survival curves were compared using the log-rank Mantel–Cox test. Data were assumed to be normally distributed. Animals were randomly assigned to experimental groups, and investigators were blinded to group allocation during experimental procedures and outcome assessment. Data are presented as mean ± SD, and n denotes the number of independent biological replicates. P < 0.05 was considered statistically significant.

## Data Availability

All data supporting the findings of this study are included in the paper and its supplementary files. Raw sequencing data reported in this study have been deposited in the Genome Sequence Archive for Human (GSA-Human) under accession HRA016454. Public datasets used in this study include GEO GSE232789, TCGA-PRAD and experiment 299759881 from the Allen Mouse Brain Connectivity Atlas. GO gene sets were obtained from Mouse MSigDB (v2026.1.Mm, M5 collection) under its standard terms of use.

## Acknowledgements

This work was sponsored by National Natural Science Foundation of China (No. 82373048); Shanghai Municipal Health Commission: Shanghai Medical Garden New Star (No. YYXX202401) - Outstanding Young Medical Talents and Excellent Projects (No. 20244Z0008); Shanghai Oriental Talent Program for Young Scholars from the Shanghai Municipal Bureau of Talent and Shanghai Municipal Health Commission (No. DFYCQN-2024-WHF); the clinical application of irreversible electroporation ablation in the treatment of metastatic prostate cancer at the Development Center of Shanghai Shenkang Hospital (No.SHDC12023107); Pudong Health Commission Leading Talent Training Program from the Pudong New Area Health Commission (No. PWRI2025-01); Specialized Prostate Center, East Hospital Affiliated to Tongji University (No. 2024-DFTS-005); Clinical and Basic Research on Prostate Cancer at Dongfang Hospital Affiliated to School of Medicine of Tongji University (No. DFRC2020006). The funders had no role in study design, data collection and analysis, decision to publish or preparation of the manuscript.

## Author contributions

H.-F.W., Y.-J.W. and W.R. contributed to the conceptualization and design of the study. W.R., P.- X.W. and Z.-Y.Q. performed neuronal tracing, chemogenetic interventions and tissue staining explant culture experiments. W.R., P.-X.W., Z.-Y.Q., S.-D.W. and R.-C.Y. established the prostate tumour models. W.R., Q.-W.Y. and J.-J.Z. performed MPG explant–tumour cell co-culture experiments. W.R. was responsible for laser-capture microdissection, Smart-seq2 and single-cell RNA sequencing-related work, transcriptomic analysis and public dataset analysis. P.-X.W., Z.-C.Y., Y.-J.S., Y.-B.W. and S.-G.C. contributed to patient recruitment, clinical specimen and information collection, and MRI data acquisition and analysis. W.R. and B.-M.H. performed image quantification, statistical analysis, data curation and figure preparation. W.R., Y.-J.W. and H.-F.W. contributed to manuscript drafting, interpretation of the results and critical review of the manuscript. W.R., P.-X.W. and Z.-Y.Q. contributed equally to this work.

---

**Extended Data Fig. 1.**
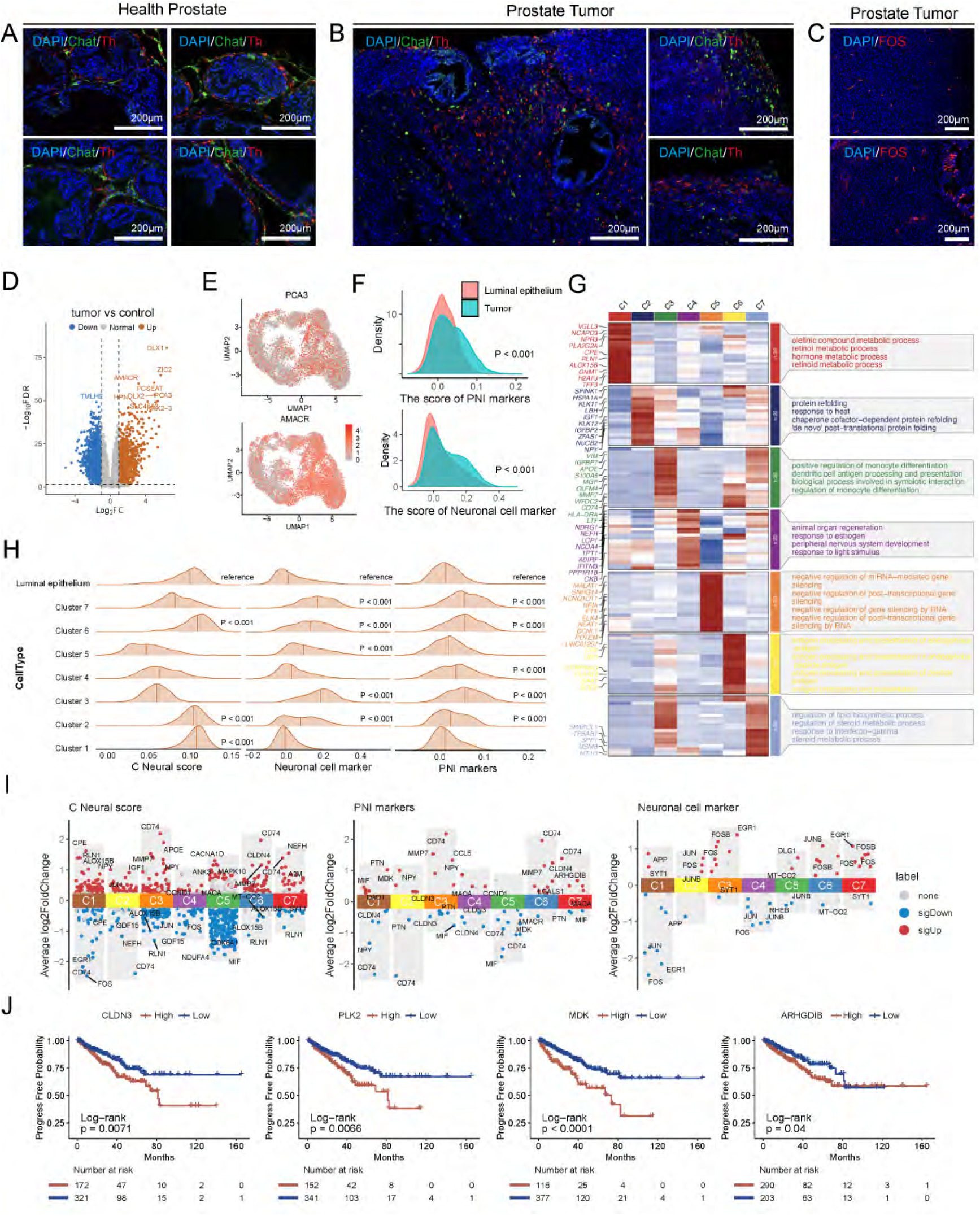
Neural infiltration and neural-like molecular features in prostate tumor. **A**, Representative immunofluorescence images showing ChAT⁺ cholinergic nerve fibers (green) and TH⁺ sympathetic nerve fibers (red) in healthy prostate tissues. **B,** Representative immunofluorescence images showing ChAT⁺ cholinergic and TH⁺ sympathetic nerve fibers in prostate tumors, with prominent neural infiltration at the tumor margins. **C,** Representative immunofluorescence images showing tdTomato⁺ nerve fibers derived from Fos-expressing neurons in prostate tumors from Ai14/FOS-CreERT2 mice. **D**, Volcano plot showing differentially expressed genes between prostate tumors and matched adjacent normal tissues in the TCGA prostate adenocarcinoma (TCGA-PRAD) cohort (|log2FC| > 1, FDR < 0.05). Red and blue indicate genes significantly upregulated and downregulated in tumors, respectively. **E,** UMAP feature plots showing the expression of PCA3 and AMACR in tumor cells. **F,** Ridge plots comparing perineural invasion (PNI) marker and neural marker gene-set scores between normal luminal epithelial cells and prostate tumor cells. Tumor cells showed significant enrichment of neural-associated molecular features. Statistical significance was assessed using Student’s t-test. **G**, Heatmap showing cluster-defining genes and representative enriched pathways in tumor-cell clusters C1–C7. **H,** Ridge plots showing PNI marker, neural marker and composite neural scores across tumor-cell clusters. Each tumor-cell cluster (C1–C7) was compared with normal luminal epithelial cells using Student’s t-test. **I,** Waterfall plots showing differentially expressed genes within the PNI marker, neural marker and composite neural-score gene sets in each tumor-cell cluster (|log2FC| > 0.25, FDR < 0.05). Red and blue indicate significantly upregulated and downregulated genes, respectively. **J,** Kaplan–Meier analysis of biochemical recurrence-free survival in the TCGA-PRAD cohort stratified by the expression of CLDN3, PLK2, MDK or ARHGDIB. High expression of each gene was associated with an increased risk of biochemical recurrence.

**Extended Data Fig. 2.**
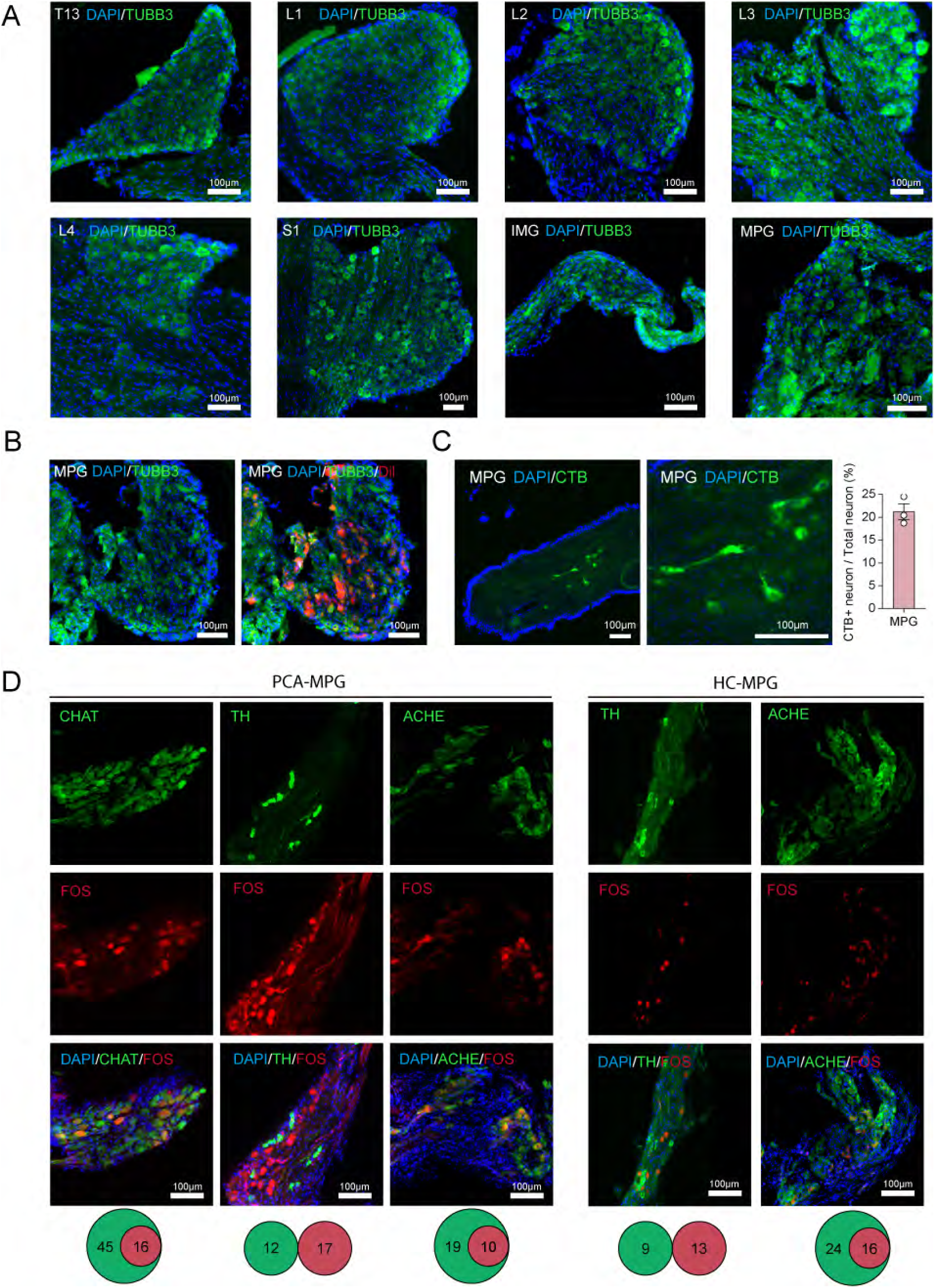
Prostate tumor-activated MPG neurons are cholinergic. **A**, Representative immunofluorescence images showing TUBB3 expression in the T13, L1, L2, L3, L4 and S1 DRGs, IMG and MPG. **B,** Representative immunofluorescence images showing colocalization of Dil-labelled prostate tumor-innervating neurons with the neuronal marker TUBB3 in the MPG of tumor mice. **C,** Representative fluorescence images showing CTB-labelled prostate tumor-innervating neurons in the MPG following retrograde tracing, together with quantification of CTB⁺ neurons as a percentage of total neurons in the MPG. n = 3 mice. **D**, Representative immunofluorescence images showing cholinergic and sympathetic neuronal markers in FOS⁺ neurons in the PCa-MPG and HC-MPG groups. In PCa-MPG, FOS⁺ neurons prominently colocalized with ChAT and AChE, but not with TH. Venn diagrams summarize the overlap between FOS⁺ neurons and the indicated markers. PCa-MPG, major pelvic ganglion from prostate tumor-bearing mice; HC-MPG, major pelvic ganglion from healthy control mice.

**Extended Data Fig. 3.**
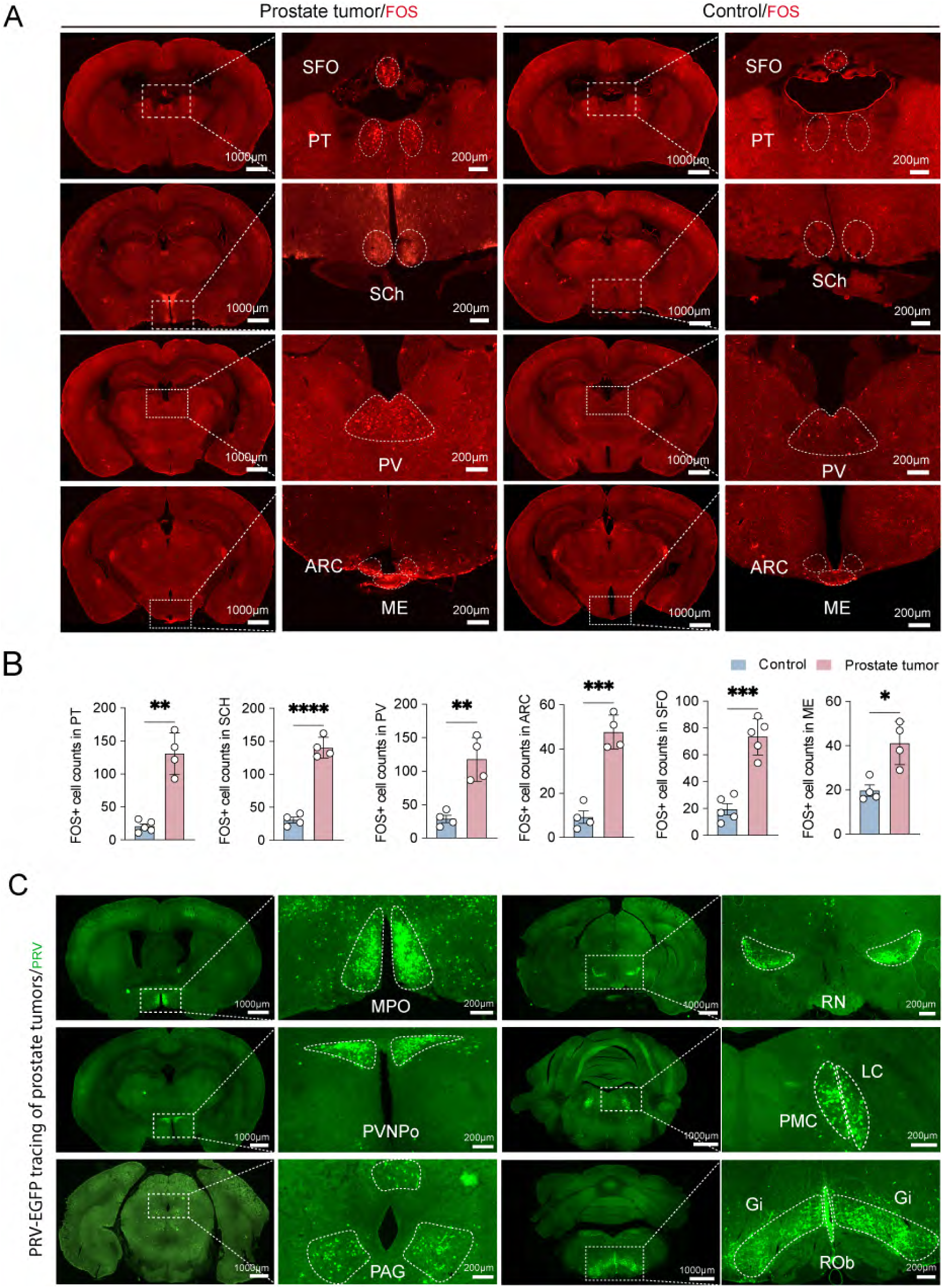
Brain region activation and neural connectivity in prostate tumor. **A**, Representative fluorescence images showing FOS activation signals in the PT, SFO, SCh, PV, ARC and ME of prostate tumor-bearing and control mice. Enlarged images of the indicated brain regions are shown alongside the corresponding whole-brain sections. PT, parataenial nucleus; SFO, subfornical organ; SCh, suprachiasmatic nucleus; PV, paraventricular nucleus of the thalamus; ARC, arcuate nucleus; ME, median eminence. **B,** Quantification of FOS⁺ cells in the indicated brain regions of control and prostate tumor-bearing mice. The numbers of FOS⁺ cells were significantly increased in prostate tumor-bearing mice compared with controls. n = 4 mice per group. Two-tailed unpaired Student’s t-test. *P < 0.05, **P < 0.01; ***P < 0.001; ****P < 0.0001. **C,** Representative fluorescence images showing PRV-EGFP-labelled neurons in multiple brain regions following trans-synaptic retrograde tracing from prostate, including the MPO, PVN, RN, PAG, PMC, LC, Gi and ROb. Enlarged images of the indicated regions are shown alongside the corresponding brain sections. MPO, medial preoptic area; RN, red nucleus; PAG, periaqueductal gray; PMC, pontine micturition center; LC, locus coeruleus; Gi, gigantocellular reticular nucleus; ROb, raphe obscurus nucleus.

**Extended Data Fig. 4.**
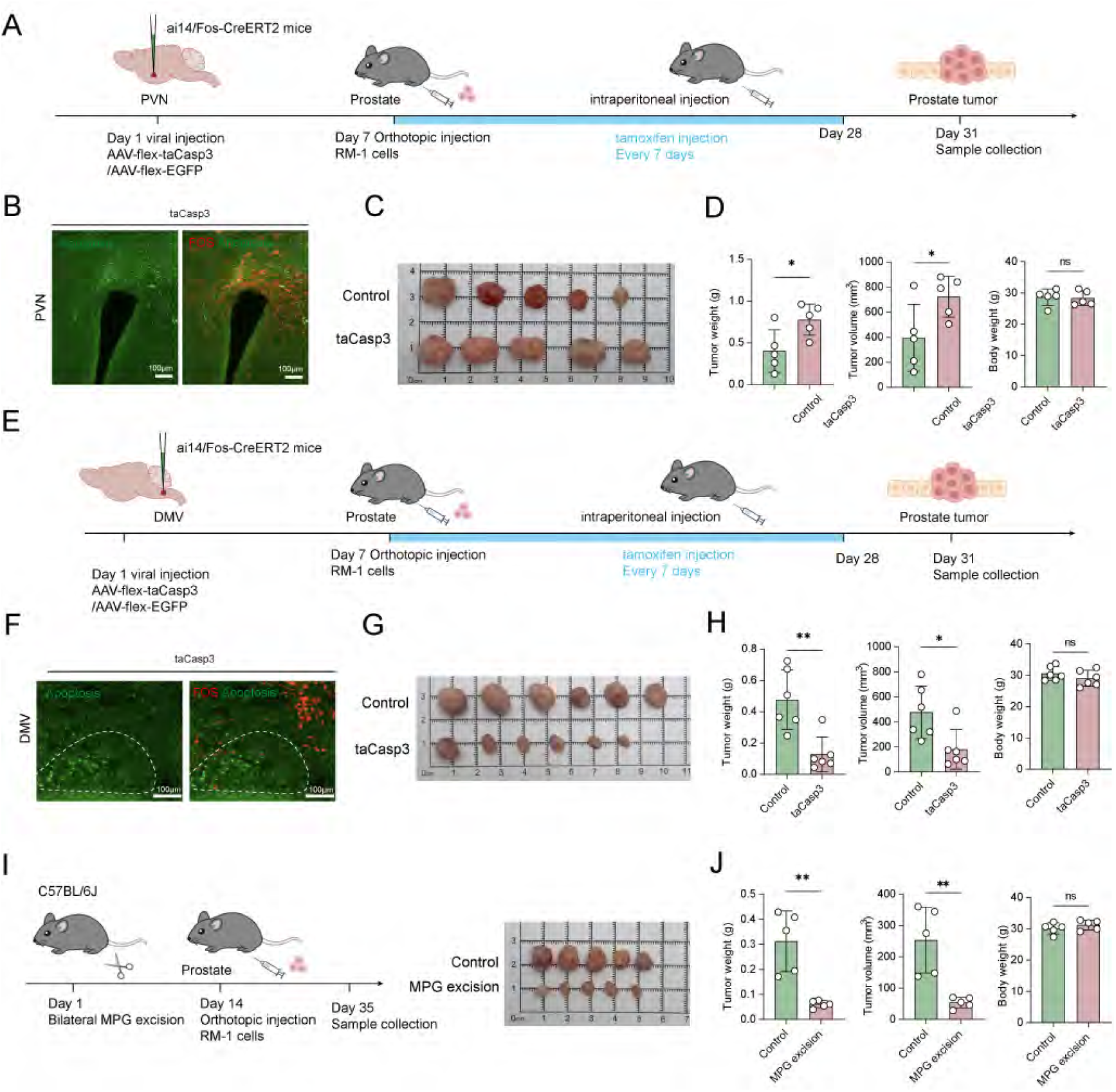
Apoptosis of activated neurons in the PVN and DMV and MPG excision affect prostate tumor growth. **A**, Experimental scheme for selective ablation of tumor-activated PVN neurons. Ai14/FOS-CreERT2 mice received PVN injections of AAV-flex-taCasp3 or control AAV-flex-EGFP. Seven days later, RM-1 cells were orthotopically implanted into the prostate, followed by tamoxifen administration according to the indicated schedule. Tumors were collected on day 31. **B–D,** Selective ablation of tumor-activated PVN neurons. Representative fluorescence images showing apoptosis in FOS-activated PVN neurons following AAV-flex-taCasp3-mediated ablation (B), macroscopic images of prostate tumors (C) and quantification of tumor weight, tumor volume and body weight (D). PVN neuronal ablation increased tumor weight and volume without affecting body weight. n = 5 mice per group. **E,** Experimental scheme for selective ablation of tumor-activated DMV neurons. Ai14/FOS-CreERT2 mice received DMV injections of AAV-flex-taCasp3 or control AAV-flex-EGFP, followed by orthotopic implantation of RM-1 cells and tamoxifen administration according to the indicated schedule. Tumors were collected on day 31. **F–H,** Selective ablation of tumor-activated DMV neurons. Representative fluorescence images showing apoptosis in FOS-activated DMV neurons following AAV-flex-taCasp3-mediated ablation (F), macroscopic images of prostate tumors (G) and quantification of tumor weight, tumor volume and body weight (H). Dashed lines delineate the DMV. DMV neuronal ablation reduced tumor weight and volume without affecting body weight. n = 6 mice per group. **I, J,** Bilateral MPG excision. Mice underwent bilateral MPG excision on day 1, followed by orthotopic RM-1 implantation on day 14 and tumor collection on day 35. Representative images of prostate tumors from control and MPG-excised mice (I). Quantification of tumor weight, tumor volume and body weight (J). MPG excision reduced tumor weight and volume without affecting body weight. n = 5 mice per group. Data are presented as mean ± SD. Two-tailed unpaired Student’s t-tests were used in D, H and J. *P < 0.05, **P < 0.01; NS, not significant.

**Extended Data Fig. 5.**
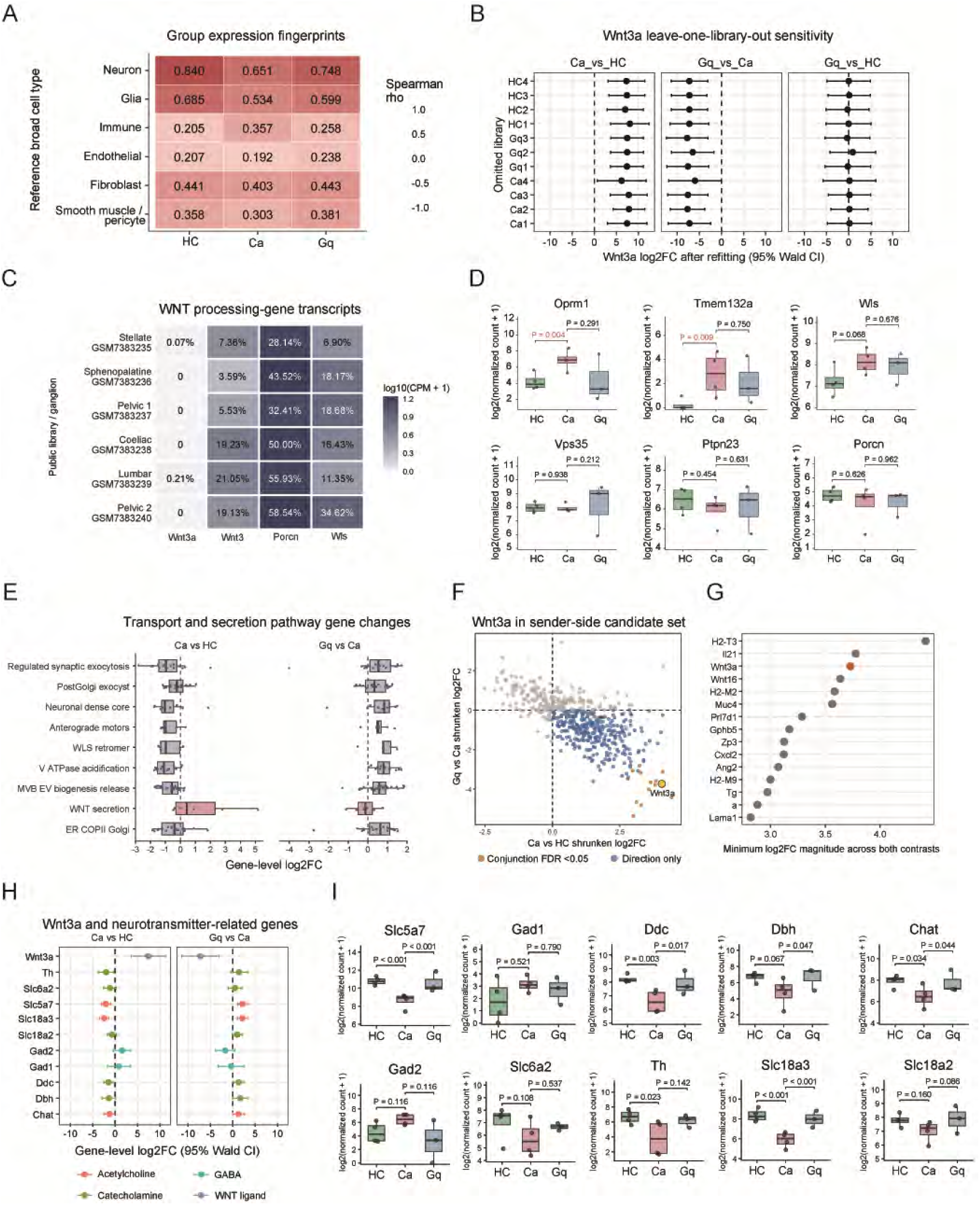
Transcriptional identity of MPG neuronal samples and expression changes in Wnt3a, transport/secretion-related genes, and neurotransmitter-related genes. **A**, Similarity between group expression profiles and public cell-type references. Reference data were obtained from mouse pelvic ganglion single-cell RNA sequencing in GSE232789, with cells assigned to six cell types based on marker genes. Pairwise comparisons between cell types, excluding activity-, injury-, and stress-related genes and Wnt3a, yielded a set of 501 identity-related genes. In-house bulk profiles were calculated as group means of log2(TPM + 1). Public single-cell data were aggregated into pseudobulk profiles by library and cell type, CPM-normalized, log2-transformed, and averaged with equal weights. Heatmap colors and values indicate Spearman correlations between the two expression profiles across the 501 genes. **B,** Leave-one-library-out sensitivity analysis of Wnt3a. One sample was omitted at a time, and differential Wnt3a expression was assessed for Ca versus HC, Gq versus Ca, and Gq versus HC. The y-axis indicates the omitted sample. Confidence intervals crossing the dashed zero line indicate no statistically significant difference. **C,** WNT-related transcripts in public peripheral ganglion neurons. Data were obtained from six mouse ganglion single-cell RNA-sequencing libraries in GSE232789. Neuronal UMIs were summed within each library, and pseudobulk CPM was calculated using the total UMIs across all genes as the denominator. Colors indicate log10(CPM + 1), and numbers within tiles indicate the percentage of neurons with at least one UMI for the indicated gene. Zero indicates no detection among the neurons included in that library. **D,** Expression of WNT protein secretion gene-set members. Expression differences across the three groups are shown for six genes in GO:0061355 (Wnt protein secretion): Oprm1, Tmem132a, Wls, Vps35, Ptpn23, and Porcn. **E,** Gene expression changes grouped by function. Each point represents the log2 fold change of a module member gene. Pink denotes the six genes in GO:0061355, and blue denotes eight functional modules. **F,** Two-contrast analysis of extracellular signaling candidate genes. The candidate set was defined by intersecting the GO extracellular space gene set with the union of GO gene sets for receptor binding, cytokine activity, growth factor activity, neuropeptide activity, or hormone activity. The directional pattern Ca > HC and Gq < Ca was assessed. Orange indicates conjunction FDR < 0.05. **G,** Ranking of directional effects among candidate genes. The x-axis indicates the smaller effect magnitude across the two expected directions of change. The 15 highest-ranking candidates showing the expected directions are displayed. **H,** Comparison of Wnt3a and neurotransmitter-related genes. Wnt3a and 10 neurotransmitter-related genes are shown, including genes involved in acetylcholine, catecholamine, and GABA synthesis, vesicular loading, or transport. Colors distinguish the indicated functional categories. Confidence intervals crossing the dashed zero line indicate no statistically significant change. **I,** Expression of neurotransmitter-related genes. Expression differences across the three groups are shown for the 10 neurotransmitter-related genes in H.

**Extended Data Fig. 6.**
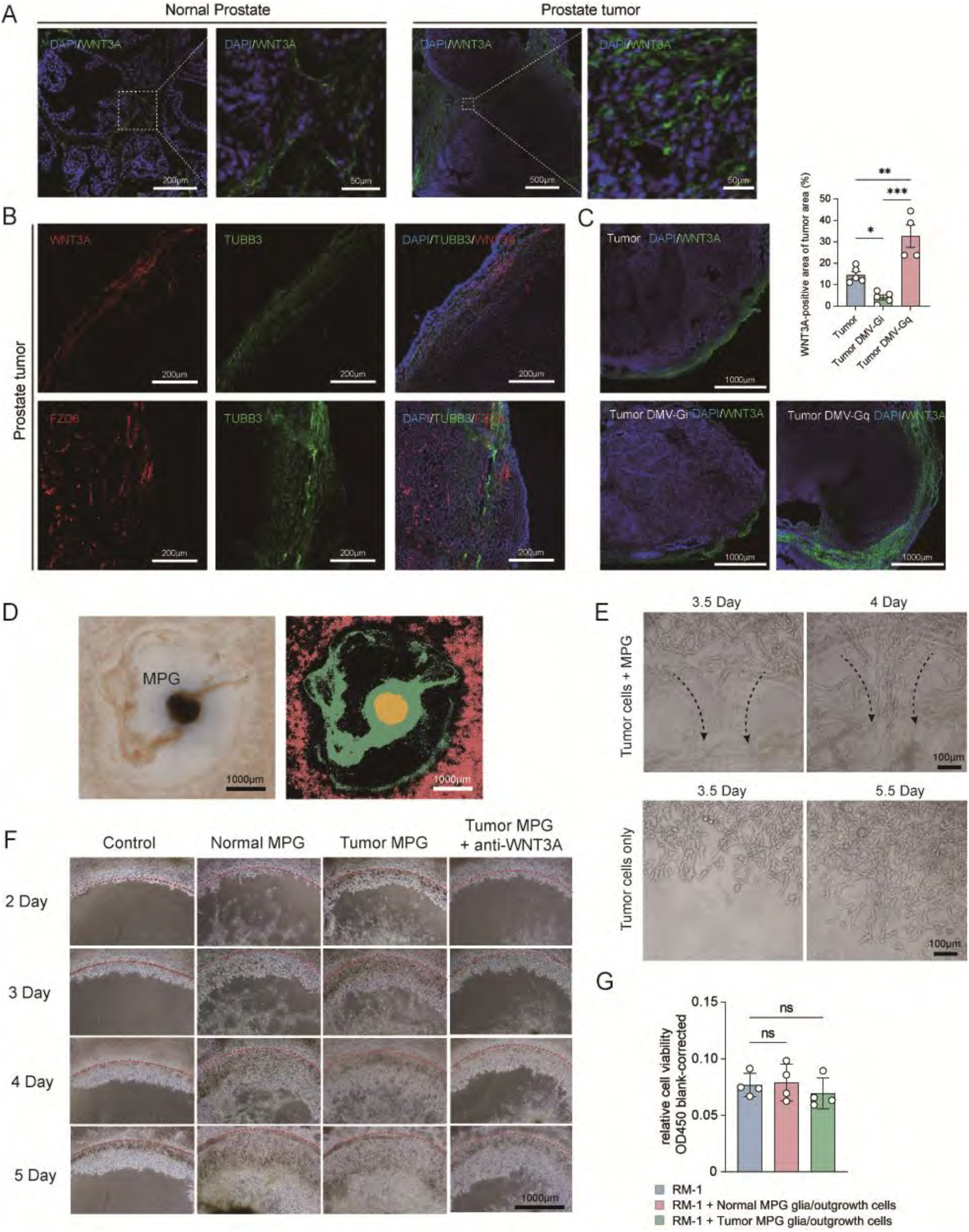
WNT3A distribution in prostate tumor tissues and co-culture of MPG explants with tumor cells. **A**, Representative immunofluorescence images showing WNT3A distribution in normal prostate and prostate tumor tissues. Enlarged images of the indicated regions are shown on the right. **B,** Representative immunofluorescence images showing the spatial distribution of WNT3A and FZD8 relative to TUBB3⁺ nerve fibers in prostate tumors. **C,** Representative immunofluorescence images and quantification of the WNT3A-positive area within prostate tumors following chemogenetic manipulation of tumor-responsive DMV neurons. Chemogenetic inhibition of tumor-responsive DMV neurons reduced the WNT3A-positive area, whereas their activation increased it. Tumor and Tumor DMV-Gi, n = 5 per group; Tumor DMV-Gq, n = 4. **D,** Representative bright-field image and corresponding pseudocolored image of the MPG explant–tumor cell co-culture system, illustrating the spatial distribution of cells surrounding the MPG explant. **E**, Representative images showing tumor cells migrating towards and aligning along nerve fibres extending from MPG explants during co-culture. Dashed arrows indicate the direction of tumor-cell migration along nerve fibres. Lower panels show tumor cells cultured without MPG, which exhibited non-directional growth. **F,** Representative images of the Matrigel invasion assay over 2–5 days for tumor cells cultured alone or co-cultured with normal MPG, tumor-associated MPG, or tumor-associated MPG in the presence of WNT3A-neutralizing antibody. Red dashed lines indicate the edge of the Matrigel. **G,** Relative viability of RM-1 cells cultured alone or co-cultured with glial cells isolated from normal or tumor-associated MPGs. Neither normal MPG-derived nor tumor-associated MPG-derived glial cells and other adherent outgrowth cells significantly altered RM-1 cell viability compared with RM-1 cells cultured alone. n = 4 per group. Data are presented as mean ± SD. One-way ANOVA followed by Tukey’s multiple-comparisons test was used in C and G. *P < 0.05, **P < 0.01; ***P < 0.001. ns, not significant.

**Extended Data Fig. 7.**
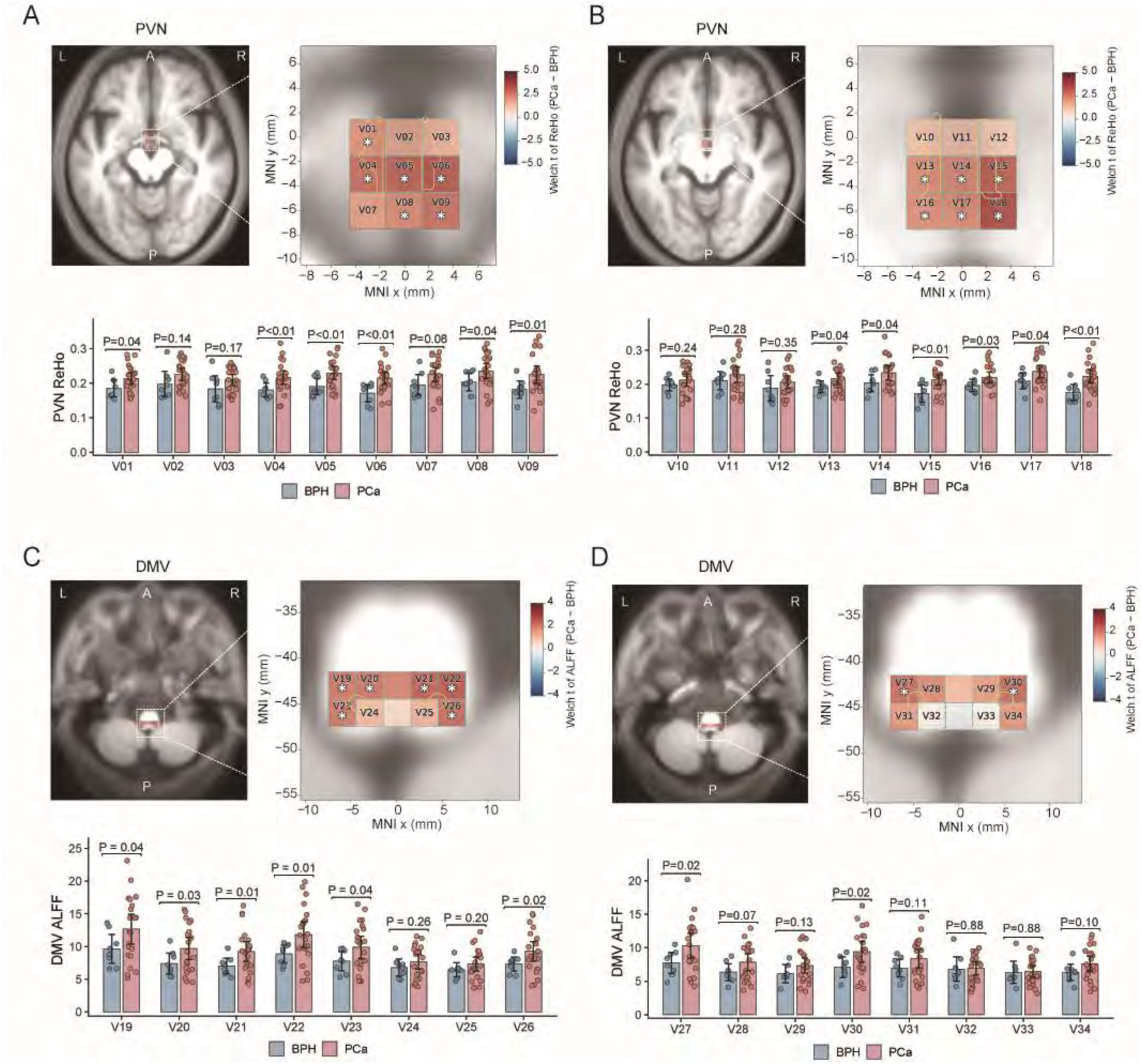
Resting-state fMRI alterations in the PVN and DMV regions in patients with prostate cancer **A, B,** Spatially smoothed regional homogeneity (sReHo) in the hypothalamic paraventricular nucleus (PVN) and adjacent regions on two consecutive axial slices at MNI z = −12 mm (A; V01–V09) and z = −9 mm (B; V10–V18). Yellow contours delineate the atlas-defined PVN. **C,D,** Spatially smoothed amplitude of low-frequency fluctuations (sALFF) in regions adjacent to the dorsal motor nucleus of the vagus (DMV) on two consecutive axial slices at MNI z = −54 mm (C; V19–V26) and z = −51 mm (D; V27–V34). Yellow DMV contours were manually drawn for anatomical illustration. Brain maps are overlaid on the mean normalized T1-weighted image across all participants. Colors indicate Welch’s t-statistics for PCa minus BPH, and asterisks indicate P < 0.05. In A and B, n = 8 for BPH and n = 22 for PCa. In the bar plots corresponding to C and D, n = 8 for BPH and n = 21 for PCa, except at V29 and V30, where BPH n = 7 owing to missing coverage. Between-group comparisons were performed using two-sided t-tests.

**Extended Data Fig. 8.**
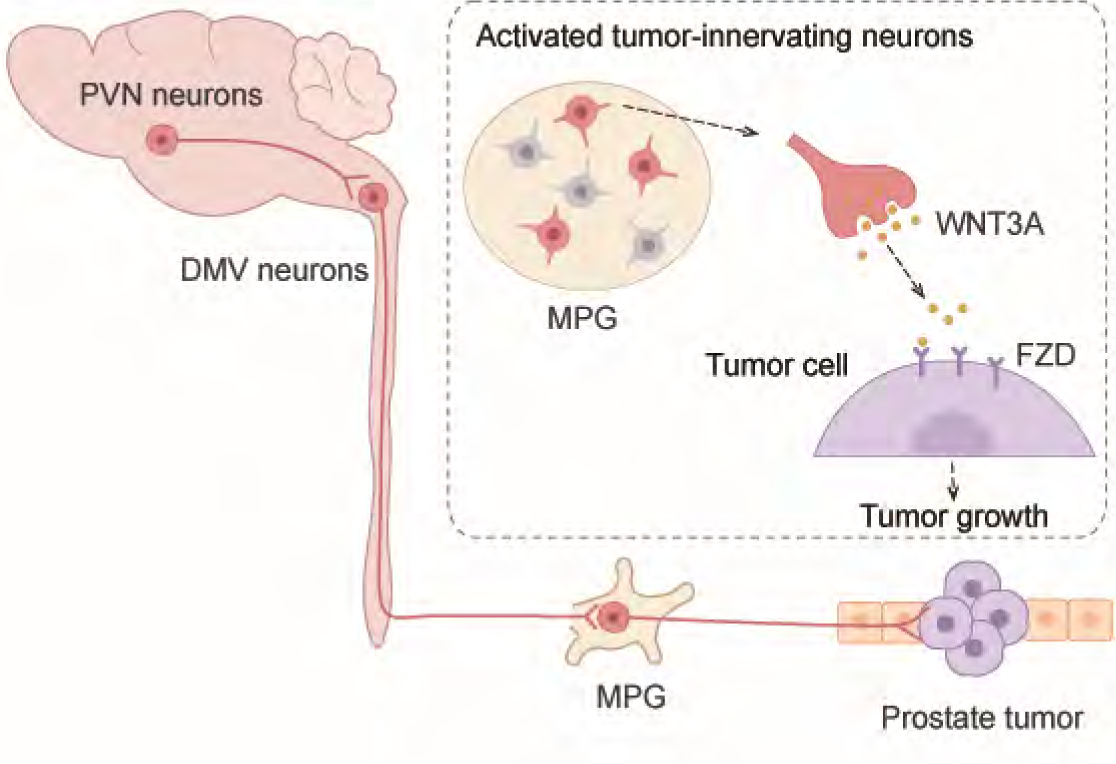
Schematic of the proposed mechanism PVN and DMV neurons are connected to tumor-innervating neurons in the major pelvic ganglion (MPG), and WNT3A derived from MPG neurons promotes tumor growth.

